# Self-supervision advances morphological profiling by unlocking powerful image representations

**DOI:** 10.1101/2023.04.28.538691

**Authors:** Vladislav Kim, Nikolaos Adaloglou, Marc Osterland, Flavio M. Morelli, Marah Halawa, Tim König, David Gnutt, Paula A. Marin Zapata

## Abstract

Cell Painting is an image-based assay that offers valuable insights into drug mechanisms of action and off-target effects. However, traditional feature extraction tools such as CellProfiler are computationally intensive and require frequent parameter adjustments. Inspired by recent advances in AI, we trained self-supervised learning (SSL) models DINO, MAE, and SimCLR on subsets of the JUMP-CP dataset to obtain powerful image representations for Cell Painting. We assessed the reproducibility and biological relevance of SSL features and uncovered the critical factors influencing model performance, such as training set composition and domain-specific normalization techniques. Our best model (DINO) surpassed CellProfiler in drug target and gene family classification, significantly reducing computational time and costs. All SSL models showed remarkable generalizability without fine-tuning, outperforming CellProfiler on an unseen dataset of genetic perturbations. Our study demonstrates the effectiveness of SSL methods for morphological profiling, suggesting promising research directions for improving the analysis of related image modalities.

## Introduction

Morphological profiling uses image-based readouts to characterize the effect of chemical and genetic perturbations^1–4^ based on alterations in cell morphology. Offering high throughput and low cost, this technology has numerous applications in drug discovery such as mode of action identification^5,6^, off-target effect detection^7,8^, drug repurposing^9–11^ and toxicity prediction^12^. One widely used assay for morphological profiling is Cell Painting that utilizes 5 fluorescent dyes to stain 8 cellular compartments^13^, generating thousands of morphological measurements per cell through automated image analysis. These high-dimensional readouts are used for hypothesis-free compound profiling, differentiating Cell Painting from target-based approaches. Despite significant progress in the visual representation learning field^14–17^, the analysis of Cell Painting images still largely relies on classical computer vision techniques^18^.

Conventional morphological profiling starts with single-cell segmentation, using CellProfiler^18^ or similar software tools^19,20^. The segmented cells are characterized using hand-crafted descriptors such as shape, size, intensity and texture^21^ among others. The descriptors are then aggregated to obtain a single vector of morphological features for each probed condition and feature selection methods are applied to reduce redundancy^21^. This multi-step workflow is computationally intensive and often requires adjustment of segmentation parameters when applied to new datasets. By contrast, deep learning models can offer a computationally efficient and segmentation-free alternative to morphological profiling.

The limited availability of biological labels has largely restricted the application of supervised learning in Cell Painting. Instead, morphological profiles are used for construction of biological maps^22,23^ to identify phenotypes and modes of action using the guilt-by-association principle. Specifically, clustering compounds or genes by morphological similarity provides mechanistic insights from annotated cluster members. Alternatively, classifiers trained on extracted features can predict downstream tasks, such as drug toxicity^24^ or cell health phenotypes^25^. However, label scarcity generally precludes end-to-end supervised learning from images, with only few exceptions^26^. The recently released JUMP-CP dataset^27^ provides unprecedented opportunities for developing novel AI-based feature extraction methods. This large-scale image set (115 TB) contains approximately 117,000 chemical and 20,000 genetic perturbations. However, most compounds lack annotations, with only 4.5% having experimentally elucidated bioactivity. Thus, leveraging the full potential of this dataset requires techniques that do not rely on data curation or biological annotations.

Self-supervised learning (SSL) methods learn feature representations from unlabeled data through a pretext task. Early SSL pretext tasks focused on predicting image transformations^28,29^. However, the current state-of-the-art performance has been achieved through methods that maximize the agreement between transformed views of the same image. For instance, PIRL^30^, MoCo^31^ and SimCLR^15^ use a contrastive loss to match paired views (“positives”) from the same image and repel unrelated views (“negatives”) from different images in the representation space. However, for optimal performance, contrastive methods require large minibatch sizes or memory banks, which can be computationally demanding. This limitation was overcome by recent non-contrastive approaches such as BYOL^32^ and DINO^16^. These approaches train a student network to predict the output of a teacher network while receiving different augmented views of an input image. Remarkably, DINO has been one of the best-performing SSL approaches across different domains^16,33,34^.

Recent advances in the SSL field have been accelerated by the adoption of the vision transformer^35^ (ViT) architecture. ViTs operate on image patches projected into tokens and use self-attention^36^ to capture global and local relationships between patches. The high computational cost of training ViT architectures inspired novel reconstruction-based pretext tasks, such as image masking, which provides a strong supervisory signal and improves training efficiency. This has been demonstrated by masked autoencoders^17^ and masked Siamese networks^37^, which achieved state-of-the-art results on natural images. Notably, ViT performance scales favorably with data volume and complexity^38^, making these models well-suited for the analysis of high-throughput imaging data.

Prior work has explored various approaches for single-cell feature extraction from high-content images, including transfer learning^39^, image inpainting^40^, variational autoencoders^41^, supervised^42^, self-supervised^23,43,44^ and weakly supervised learning^45–47^. However, existing single-cell methods require segmentation, leading to complex multistep workflows. Other approaches extract representations from whole images ^48–51^, but rely on scarce labels or pretrained ImageNet weights, which restricts their application to 3-channel images or requires embedding concatenation for multichannel (> 3) applications. To date, only few approaches^52,53^ learn directly from microscopy images without segmentation or manually curated annotations. Moreover, a systematic study evaluating the benefits of SSL methods over classical analysis workflows for high-content imaging data is currently missing.

Here, we present the first comprehensive benchmark study of state-of-the-art SSL methods adapted for Cell Painting images. We trained all SSL models directly in the applicability domain and assessed their generalizability on independent datasets with chemical and genetic perturbations. Crucially, we examined the requirements for the successful application of SSL in the fluorescence microscopy domain including relevant image augmentations, model architecture, training set composition and feature postprocessing techniques. We assessed the performance gap between supervised and SSL models for compound bioactivity prediction, elucidating scenarios favoring each approach. Our results indicate that SSL methods provide a robust, efficient, and segmentation-free alternative to CellProfiler and that SSL features enable accurate prediction of compound properties comparable to supervised models.

## Results

### SSL framework for segmentation-free morphological profiling

Our self-supervised learning (SSL) framework operates directly on image crops without cell segmentation (see Methods). We adapted 3 state-of-the-art SSL approaches (**Fig. 1a**) for 5-channel Cell Painting images: SimCLR (simple framework for contrastive learning of visual representations)^15^, DINO (distillation with no labels)^16^ and MAE (masked autoencoder)^17^. We pretrained all SSL methods on two JUMP-CP^27^ data subsets: single-source and multisource training sets (see Methods). Using these pretrained models, we extracted features to construct morphological profiles of chemical and genetic perturbations in held-out evaluation sets (**Fig. 1c**). We benchmarked the SSL features against two baselines (**Fig. 1b**): CellProfiler^18^, a widely used computational tool for morphological profiling, and transfer learning from a model pretrained on natural images^35^ (see Methods).

**Figure 1:**
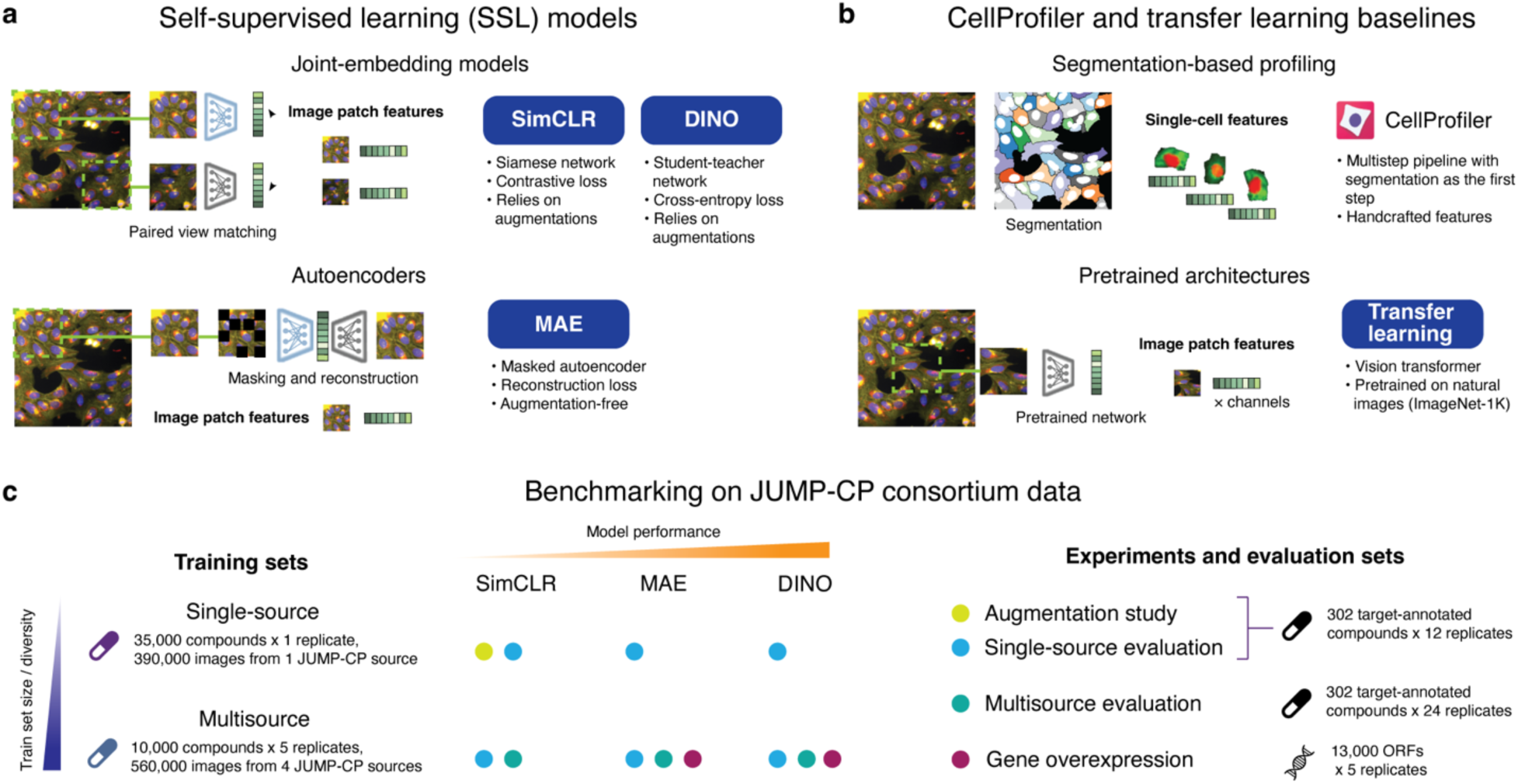
Self-supervised learning for morphological profiling. **a)** Schematic of the SSL models used in this study. All models are segmentation-free and use only image crops as input. SimCLR and DINO were trained on the pretext task of matching features from augmented views of the same image. MAE was trained on the image reconstruction task with partially masked input. **b)** Schematic of the two baseline methods. CellProfiler: conventional method based on single-cell segmentation and handcrafted features. Transfer learning: pretrained vision transformer that outputs per-channel features. **c)** JUMP-CP consortium data subsets used for training and evaluation of SSL models. SSL models were trained on two training sets: a single-source and a multisource set. The colored points indicate which evaluation sets were used to assess the models trained on the single-source and multisource data. The image augmentation study was conducted only for SimCLR trained and evaluated on the single-source data.

We used small (ViT-S) and base (ViT-B) vision transformer architectures as SSL backbones that encode images into feature vectors. At inference, we split images into equally-sized crops that were input into a pretrained ViT, with the image feature obtained by averaging the crop features (see Methods). To correct for plate and experimental batch effects, we tested several normalization methods and selected the best postprocessing strategy for each feature type (see Methods and **Extended Data Fig. 1-2**). We generated perturbation profiles by averaging normalized features across replicates of the same perturbation. As JUMP-CP datasets contain several data sources and experimental batches (see Methods), this aggregation was conducted at the batch, source, and full dataset levels (see Methods), enabling assessment of reproducibility across batches and sources. Full dataset aggregation produced consensus profiles, which were used for drug target and gene family classification.

### Benchmarking feature extraction methods on JUMP-CP data

We evaluated all feature extraction methods on 3 held-out JUMP-CP^27^ data subsets (**Fig. 1c**). The first two subsets contained target-annotated compounds with 2 drugs per target class (see Methods). This allowed us to assess the suitability of pretrained features for few-shot learning, an important task in morphological profiling, where only few examples per class are available. The third evaluation set consisted of gene overexpression perturbations (see Methods) from a data source not used in training. Including genetic perturbations allowed us to test the models’ ability to generalize to previously unseen perturbations, since our SSL models were trained only on images of chemically perturbed cells. As all JUMP-CP perturbations were screened across multiple experimental batches in different laboratories (see Methods), we could also assess batch and data source effects.

We compared feature extractors based on two key criteria: reproducibility and biological relevance (see Methods). To assess reproducibility, we used *perturbation mAP* (*mAP:* mean average precision), which quantifies the agreement between replicates of the same perturbation across experimental batches. Biological relevance was evaluated by the agreement between perturbations with the same biological annotation, using *target mAP* as the metric (see Methods). Additionally, we used nearest neighbor (NN) accuracy of matching across experimental batches (*not-same-batch* or *NSB accuracy*) and across experimental batches and distinct perturbations (*not-same-batch-or-perturbation* or *NSBP accuracy*). For genetic perturbations, we additionally evaluated the clustering quality with respect to gene family labels, using the *adjusted mutual information* (*AMI*) metric (see Methods).

### Establishing augmentations for Cell Painting images

SimCLR and DINO rely on image augmentations to generate views of the same image during training (Fig. 2a). While the relative importance of individual augmentations was previously determined for RGB images^15^, there are no prior works systematically assessing augmentations for multichannel microscopy images. Therefore, we conducted a comprehensive study for Cell Painting images to evaluate the contribution of several common augmentations. We followed a similar study design as in ^15^, probing all pairwise combinations of 5 augmentations (Fig. 2a): ‘Resize’, ‘Color’, ‘Drop channel’, ‘Gaussian noise’ and ‘Gaussian blur’ (see Methods). Since color jittering^15,16^ is specific to RGB images, we replaced the ‘Color’ augmentation with a more general transform consisting of random brightness change and intensity shift applied to each fluorescent channel independently (see Methods).

**Figure 2:**
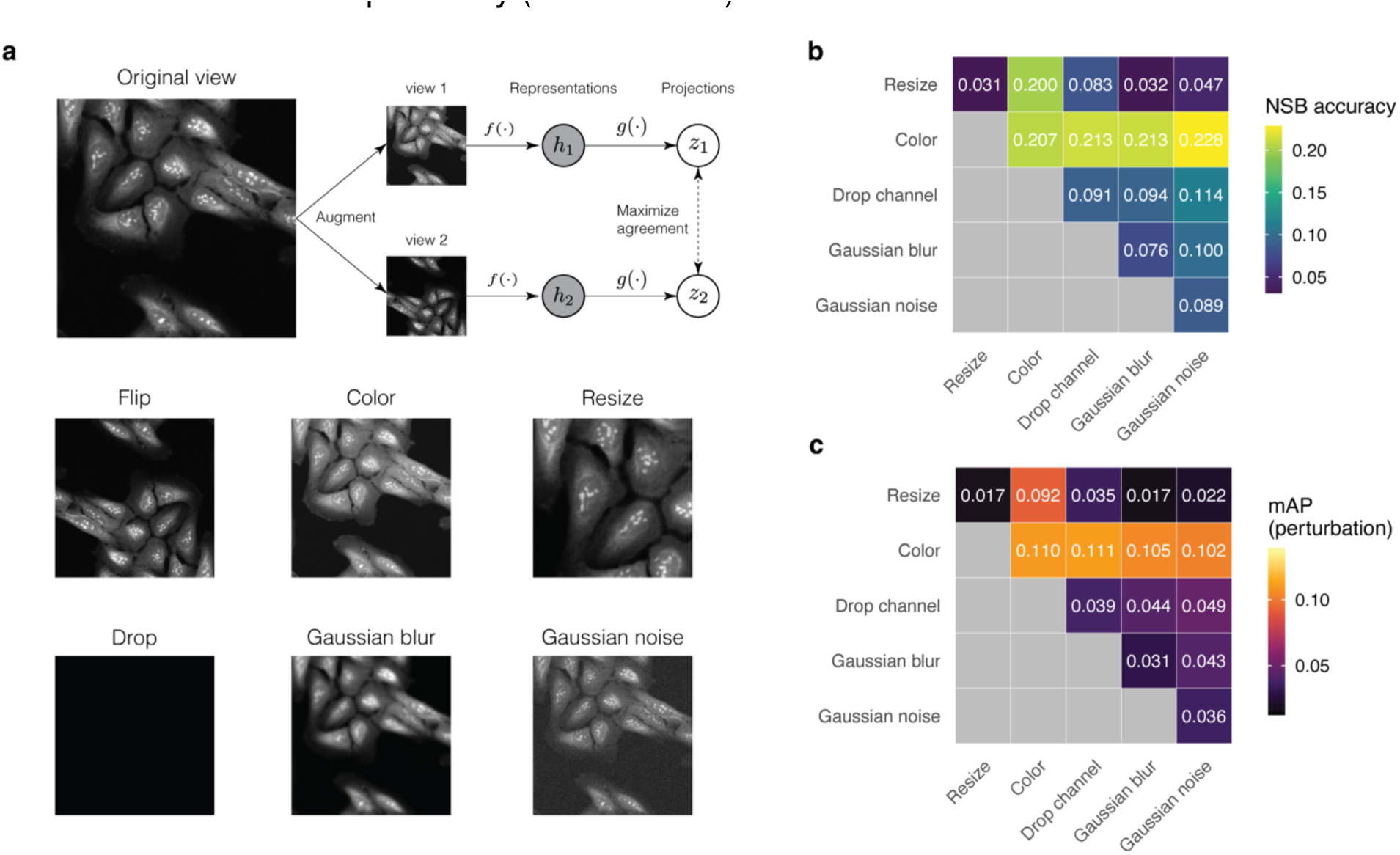
Augmentations for multichannel microscopy images. **a)** Schematic of the SimCLR contrastive learning framework and representative images of the tested augmentations. Flip: vertical and horizontal flips. Color: per-channel stochastic intensity shift and brightness adjustment. Resize: random crop of variable dimensions followed by rescaling to a fixed crop size. Drop: omit one color channel at random. Gaussian blur: Gaussian kernel smoothing. Gaussian noise: addition of Gaussian noise. For details, refer to the Methods section. **b)-c)** Performance comparison of 15 SimCLR models with different combinations of augmentations trained and evaluated on the single-source data. Diagonal entries report the performance of individual augmentations. The ‘Flip’ augmentation was applied by default. Performance is assessed based on the reproducibility metrics *not-same-batch accuracy* (*NSB*) and *perturbation mean average precision* (*mAP*) (see Methods).

We pretrained 15 SimCLR models with different augmentation strategies (see Methods). A pairwise comparison of these models (Fig. 2b**-c**) revealed that the ‘Color’ augmentation had the greatest positive impact on performance both in terms of *perturbation mAP* and *NSB accuracy*. The ‘Resize’ operation, which generates image crops at different scales (see Methods), led to a decrease in performance compared to other augmentations. The remaining 3 augmentations (‘Drop channel’, ‘Gaussian noise’, ‘Gaussian blur’) had a negligible effect on *NSB accuracy* and *mAP* relative to the ‘Color’ augmentation alone. Based on these findings, we established an augmentation pipeline for Cell Painting images that relies primarily on ‘Color’ and ‘Flip’ augmentations and excluded all other transforms which didn’t improve performance. This pipeline was adopted for DINO and SimCLR in all subsequent experiments.

### DINO trained on multisource data outperforms CellProfiler

First, we evaluated the performance of DINO, MAE and SimCLR trained on different datasets (Fig. 3a**,b**). While all SSL models trained on single-source data performed worse than CellProfiler, models trained on multisource data match (MAE) or exceed (DINO) CellProfiler performance (Fig. 3a**,b**). DINO trained on multisource data achieved the best results among the SSL methods, surpassing CellProfiler. DINO features displayed better reproducibility (Fig. 3a**,c**) and biological relevance (Fig. 3b**,d**) compared to MAE and SimCLR. On the first evaluation set (Fig. 3a**-b**), DINO surpassed CellProfiler by a margin of 16% in *perturbation mAP*, 3% in *NSB accuracy,* 22% in *target mAP,* and 32% in *NSBP accuracy*. On the second evaluation set (Fig. 3c**-d**), DINO outperformed CellProfiler by even greater margins: 29% in *perturbation mAP*, 6% in *NSB accuracy*, 61% in *target mAP*, and 11% in *NSBP accuracy*. Notably, transfer learning features showed the worst performance, encouraging the use of SSL methods for morphological profiling.

**Figure 3:**
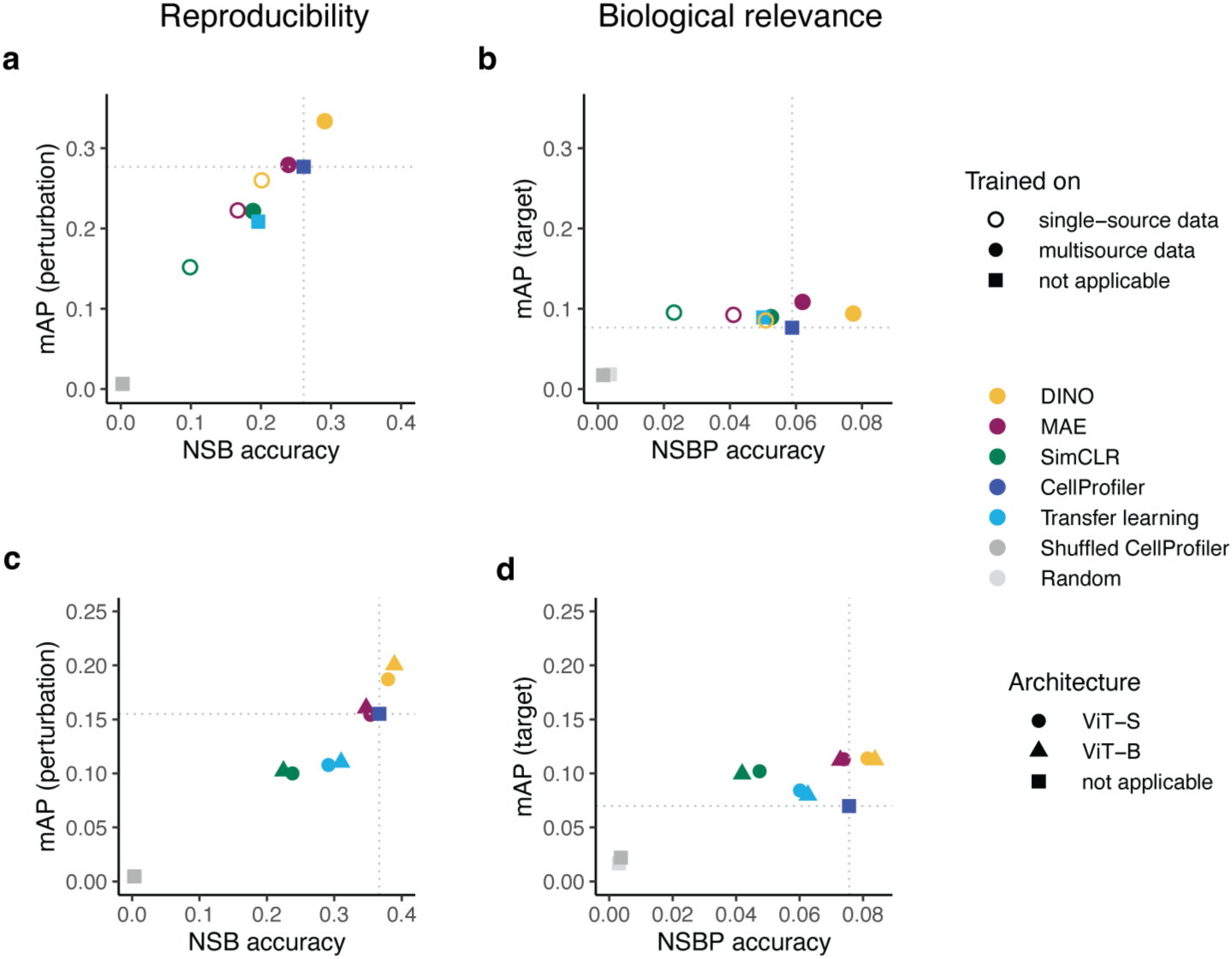
Impact of training data and model size on SSL model performance. Performance comparison of SSL models (DINO, MAE, and SimCLR) and two baselines (CellProfiler and transfer learning) for different datasets (single-source and multisource) and model architectures (ViT-S and ViT-B). Left panels: reproducibility metrics *perturbation not-same-batch (NSB) accuracy* and *mean average precision (mAP).* Right panels: biological relevance metrics *target not-same-batch-or-perturbation (NSBP) accuracy* and *mean average precision (mAP)*. Colors indicate different models and two randomized baselines: Shuffled CellProfiler (CellProfiler features with shuffled labels) and Random (random normally distributed features). Dotted lines indicate CellProfiler performance. **a)-b)** Performance on the single-source evaluation set. Shapes indicate different training sets. Only the results for ViT-S architectures are reported. **c)-d)** Performance on the multisource evaluation set. Shapes indicate the ViT architecture. All SSL models were trained on the multisource training set.

The superiority of DINO was even more evident in F1-score curves (**Extended Data Fig. 3**) and further confirmed by comparing reproducibility across JUMP-CP data sources (**Extended Data Fig. 4**, see Methods). DINO yielded similar or better performance to CellProfiler in 3 out of 4 JUMP-CP data sources (**Extended Data Fig. 4a-d**) and achieved higher *perturbation mAP* (+73%) and *NSB accuracy* (+2%) on a cross-source matching task (**Extended Data Fig. 4e**, see Methods).

Interestingly, models based on the larger ViT-B architecture only showed a marginal improvement over the ViT-S models (Fig. 3c**,d**). This was observed for both the SSL methods and the transfer learning baseline. Our results indicate that the biggest performance gain was achieved by expanding the training set to a more extensive and diverse image set as opposed to increasing the model size. In subsequent evaluations, we focused solely on SSL models trained on the multisource data using the ViT-S architecture, motivated by its lower compute requirements.

### UMAP embeddings reveal biological and technical axes of variation

Next, we used UMAP^54^ (see Methods) to embed DINO, MAE, SimCLR and CellProfiler features in 2 dimensions. To assess whether feature embeddings produced biologically meaningful clusters, we highlighted a selection of 20 drug targets in the UMAP space (see Methods). All embeddings grouped compounds with the same target to some extent (Fig. 4). Notably, DINO and CellProfiler embeddings yielded well-separated clusters in the UMAP, highlighting targets such as NAMPT, PAK1, and RET (Fig. 4a, d). Additionally, DINO embeddings demonstrated superior cluster separation for KRAS, AKT1, DNMT3A, TGFBR1, CDK2, and CDK7 (Fig. 4a) compared with CellProfiler (Fig. 4d). These results further support that DINO features are biologically meaningful and at least as powerful as those of CellProfiler.

**Figure 4:**
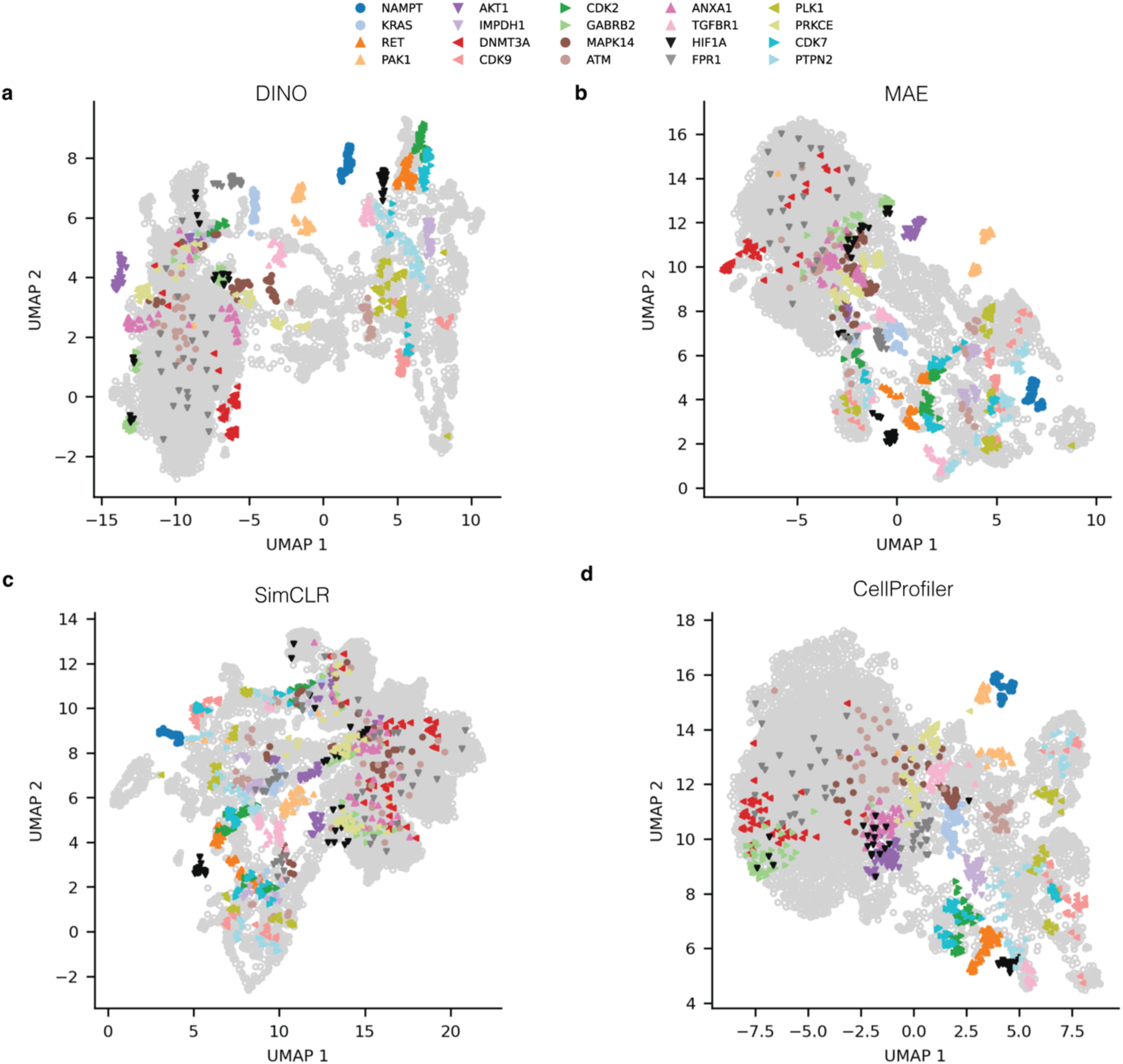
UMAP embeddings of SSL and CellProfiler features reveal compound clusters. Two-dimensional embeddings of well-level features from SSL methods and the CellProfiler baseline on the multisource evaluation set. Colors highlight target labels for a selection of 20 targets (see Methods). Perturbations with other targets are depicted as grey hollow points. All SSL models used the ViT-S architecture.

We then examined feature robustness with respect to technical sources of variation by coloring UMAP embeddings by experimental batch and data source (**Extended Data Fig. 5a,b**). SSL feature embeddings were more susceptible to technical variations compared to CellProfiler, displaying a stronger separation between experimental subgroups. Upon closer inspection, we found that the most pronounced data source effects occurred within DMSO negative controls (**Extended Data Fig. 5a,c**) and that the differences strongly correlated with variations in cell count (**Extended Data Fig. 5d**). We hypothesize that CellProfiler is more robust towards cell count variations since the features are extracted from single cells. Additionally, we quantified the impact of technical variation (see Methods) and found that both SSL and CellProfiler features were affected by experimental batch and source effects to some extent (**Extended Data Fig. 6**). These results suggest that SSL representations capture both biological and technical axes of variation in the data.

### DINO features generalize to genetic perturbations and recapitulate gene families

To evaluate the generalizability of SSL models, we used an independent dataset of gene overexpression perturbations from a new source (see Methods). As for chemical perturbations, DINO features demonstrated the best reproducibility (Fig. 5a), with MAE achieving comparable performance. We also assessed the ability to predict gene family labels (see Methods) and found that all SSL features outperformed CellProfiler on metrics of biological relevance (Fig. 5b**-c**). The most notable improvements were seen with DINO and MAE features that improved gene family predictions by 41% in *NSBP accuracy* over CellProfiler.

**Figure 5:**
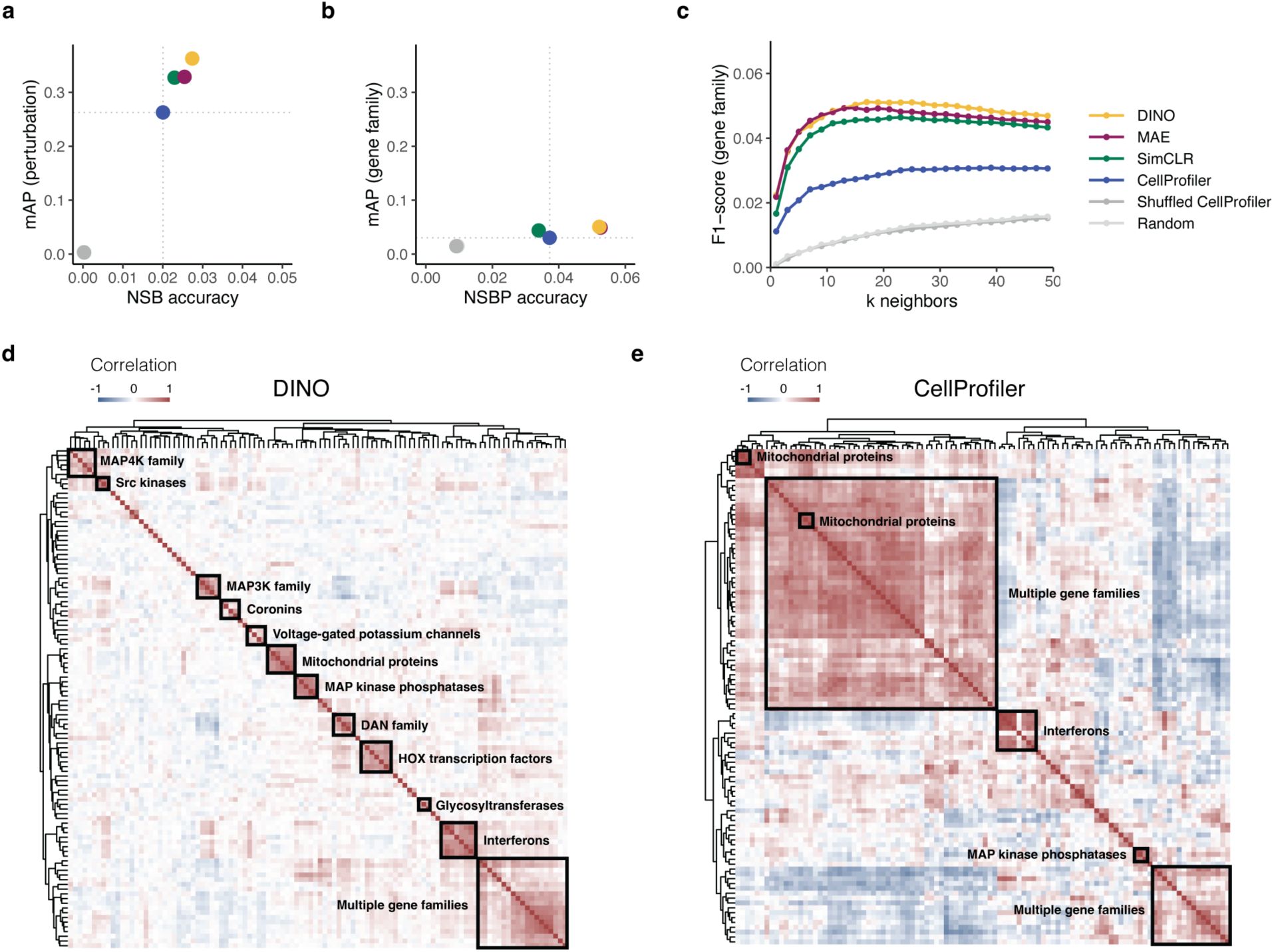
Evaluation of SSL models on genetic perturbations demonstrates generalizability to unseen data. Evaluation of SSL models and CellProfiler on an independent gene overexpression set. SSL models were trained only on images of chemically perturbed cells. Colors indicate different models and two randomized baselines: Shuffled CellProfiler (CellProfiler features with shuffled labels) and Random (random normally distributed features). Dotted lines in a)-b) indicate CellProfiler performance. **a)** Reproducibility metrics: *perturbation not-same-batch (NSB) accuracy* and *mean average precision (mAP)*. **b)** Metrics of biological relevance: *gene family not-same-batch-or-perturbation (NSBP) accuracy* and *mean average precision (mAP).* **c)** F1-scores for matching gene family labels based on gene consensus profiles for a range of nearest neighbors *k.* **d)-e)** Hierarchical clustering of the 20 gene families with the highest intragroup correlations in the DINO and CellProfiler representation spaces, respectively. Detailed versions of the heatmaps displaying gene and gene family annotations for each row are presented in **Extended Data Fig. 7-8**.

Using pairwise gene similarity analyses, we tested the ability of morphological features to recapitulate gene families. For a selection of 20 gene families (see Methods), we performed hierarchical clustering of gene profiles for DINO (Fig. 5d) and CellProfiler (Fig. 5e). The resulting similarity maps were annotated to highlight groups based on gene and gene family labels (see Methods). Our qualitative comparison (Fig. 5d **and Extended Data Fig. 7**) revealed that DINO features recovered a larger number of gene groups and produced more homogeneous clusters than CellProfiler. DINO recapitulated 11 gene groups, including MAP4K family, Src kinases, MAP3K family, coronins, voltage-gated potassium channels, mitochondrial proteins, MAP kinase phosphatases, DAN family, HOX transcription factors, glycosyltransferases, and interferons. By contrast, CellProfiler recovered only 3 gene groups (mitochondrial proteins, interferons, and MAP kinase phosphatases) and 2 large clusters of mixed gene families (Fig. 5e **and Extended Data Fig. 8**).

To provide a more objective assessment, we computed the adjusted mutual information (AMI) between cluster assignments and gene family labels (see Methods). We found that DINO (AMI = 0.51) outperformed CellProfiler (AMI = 0.19) on the gene clustering task, indicating that DINO features excel at capturing gene family information. Since all SSL models were trained on images with compound-treated cells, these results demonstrate the remarkable generalizability of SSL models, enabling their application to unseen data sources and conditions without parameter adjustments.

### DINO enables bioactivity prediction comparable to supervised CNNs

In ^26^, convolutional neural networks (CNNs) were used to predict compound activity across 209 ChEMBL assays from Cell Painting images. To evaluate the performance gap between supervised and self-supervised learning, we compared bioactivity prediction models trained on DINO features versus images directly. Specifically, we assessed a neural network (NN) trained on DINO features (see Methods) against 6 CNNs trained on Cell Painting images from ^26^. We used DINO pretrained on the JUMP-CP data, allowing us to probe its out-of-distribution generalizability. As an additional baseline, a NN trained on CellProfiler features was incorporated from ^26^.

After ranking all methods by mean AUCROC across 209 assays (see Methods), we found (**Extended Data Table 1**) that the model trained on DINO features (AUCROC = 0.72) achieved performance comparable to GapNet (AUCROC = 0.73), the third best method. Although the top 3 CNNs had slightly higher mean AUCROC values, DINO outperformed 3 additional CNNs and a model trained on CellProfiler features (**Extended Data Table 1**). Among the 8 methods compared, DINO ranked 4th for the number of assays predicted with AUCROC above 0.7 and 0.8. This confirms that DINO can generalize to novel datasets and tasks without fine-tuning.

Nevertheless, the top 3 CNNs surpassed the DINO-based model in the number of assays predicted with AUCROC > 0.9, indicating a performance gap. Further analysis revealed that the model trained on DINO features offered comparable or better performance on assays with limited data (**Extended Data Fig. 9**). For assays with only few (< 100) activity labels available, the DINO model produced a similar number of accurate predictions as the top 3 CNNs (**Extended Data Fig. 9a**). However, on assays with 100-500 labels, the CNNs showed better performance than the DINO model (**Extended Data Fig. 9a**). When considering only assay predictions with AUCROC > 0.7 (**Extended Data Fig. 9b**), the DINO model achieved higher median AUCROC values for assays with few activity labels (< 50 and 50-100), consistent with SSL features excelling in few-shot settings^55^.

### SSL pipeline is significantly faster than CellProfiler

As efficient data processing is crucial for accelerating research throughput, we additionally benchmarked the computation time and cloud costs of feature extraction using DINO versus CellProfiler. For this comparison, we used 12 GPU-accelerated cloud instances for DINO and 12 CPU-intensive instances for CellProfiler (**Extended Data Table 2**). We found that DINO was 50 times faster than CellProfiler, with an average processing time of 1.3 minutes per plate (**Extended Data Table 2**). Despite the need for GPU resources, the cloud costs per plate were over 50 times lower for DINO than for CellProfiler (**Extended Data Table 2**). Moreover, DINO offers a simpler workflow which processes images end-to-end, in contrast to the multi-step CellProfiler workflow which requires illumination correction, segmentation, feature extraction and selection.

Taken together, our findings show that image-level SSL methods are a viable alternative to traditional segmentation-based approaches, offering improved performance, generalizability to new datasets, speed, and lower workflow complexity and computational costs.

## Discussion

To assess the applicability of SSL for Cell Painting, we trained and evaluated 3 state-of-the-art methods DINO, MAE, and SimCLR on complex datasets with chemical and genetic perturbations. Using reproducibility and biological relevance as our main criteria, we showed that our best model, DINO, outperformed the established feature extraction tool, CellProfiler, in drug target and gene family classification, with even greater improvements in gene clustering.

Our SSL models captured informative cell-related features that generalized to unseen datasets without parameter fine-tuning. While trained only on compound perturbations, DINO achieved superior classification and clustering performance on a novel gene overexpression set, facilitating the construction of biological maps^22^. For compound activity prediction, DINO features transferred remarkably well to a new dataset, with the model trained on DINO features achieving comparable performance to CNNs^26^ trained on that dataset directly. We hypothesize that the strong transferability^56^ of DINO can be attributed to its image-level pretext task that effectively captures low-frequency signals^57^ associated with cellular shape. The generalizability of our SSL models expedites the analysis of new datasets in contrast to CellProfiler, which requires frequent parameter adjustments.

Previous studies^45,47,51^ fine-tuned ImageNet-pretrained networks to learn representations for Cell Painting, often curating the training set through compound preselection. These methods process each channel independently and output concatenated channel representations, increasing computational complexity and feature redundancy. By contrast, our SSL models are tailored for uncurated 5-channel images, resulting in compact representations with lower redundancy. More recently, DINO was applied for learning single-cell morphological representations^43,44^, with ^44^ reporting superior performance over CellProfiler. However, unlike these approaches, our SSL framework operates without cell segmentation, streamlining feature extraction.

Our study offers practical guidance for applying SSL methods to microscopy images. We systematically examined the role of augmentations, confirming the importance of ’Color’ augmentation observed for natural images^15^. We found, however, that the ’Resize’ augmentation decreased representation quality. We hypothesize that enforcing scale-invariance is detrimental, as cell size variations in microscopy images reflect actual phenotypic changes. Additionally, training set size and heterogeneity played a crucial role in SSL model performance. Scaling from a single-source to a multisource training set improved performance more than using larger vision transformer architectures, which may require more training data and longer pretraining^58^. Notably, masked image modelling (MAE) performed comparably to self-distillation pretraining (DINO), which suggests that combining both SSL pretext tasks could further improve representations as shown for natural images^59,60^.

Our analysis (**Extended Data Fig. 9**) revealed that models trained on SSL features excel in bioactivity prediction when ground-truth labels are scarce, while dedicated supervised methods achieve superior performance given ample labeled data. Since compound annotations are sparse, training supervised models directly from images remains infeasible in most but few^26^ cases, which makes the use of SSL features an attractive alternative to harness the large-scale unlabeled data such as the JUMP-CP dataset.

With a 50-fold reduction in compute time and costs compared to CellProfiler, SSL feature extraction methods can facilitate compound screening campaigns of unprecedented scale, revolutionizing the pace of early drug discovery. However, pretraining a DINO model demands substantial compute resources, requiring approximately 300 GPU hours. Given the intensive resource requirements, our study used only subsets of the JUMP-CP dataset, leaving room for exploration of the full dataset’s potential. Investigating the emerging properties of larger self-supervised ViTs trained on the complete JUMP-CP dataset offers promising research directions.

One limitation of our segmentation-free approach is that it operates at image-crop level and does not provide insights into cell heterogeneity, making CellProfiler a more suitable tool for single-cell analyses. Additionally, CellProfiler provides interpretability, by linking individual features to specific microscopy channels and mathematically defined morphological descriptors. However, even with self-supervised ViTs, we can gain some level of interpretability by examining self-attention maps (see **Extended Data Fig. 10**). Furthermore, SSL methods showed higher susceptibility to experimental batch and laboratory effects compared to CellProfiler. Post-hoc approaches like Harmony^61^ can mitigate the effect of technical variation on SSL features. Alternatively, incorporating batch alignment as an additional objective during pretraining may produce more robust SSL representations.

Our SSL models showed generalizability across Cell Painting datasets but remain limited in transferability across other microscopy modalities, requiring fine-tuning for assays with different staining than Cell Painting. Drawing inspiration from natural language and image domains^62–65^, we encourage the development of assay-agnostic foundation models for microscopy images, which can standardize and expedite the analysis of high-content assays across various imaging modalities.

## Methods

### Technical terminology

Image *features* are high-dimensional readouts extracted from images using segmentation-based or deep learning approaches. We use the terms “*representations*” and “*features*” interchangeably. Feature vectors can be embedded into 2 dimensions for visualization; we refer to these projections as *embeddings*.

Cell Painting assay is conducted in 384-well *plates*, with each well imaged at several locations to produce multiple images or fields of view (FOVs). *Well profiles* or *well features* refer to features aggregated for each well across multiple FOVs. Cell Painting screening is performed in experimental *batches* containing groups of plates. Unless specified otherwise, the term “batch” refers to an experimental batch and not to a minibatch used for training deep learning models. In the JUMP Cell Painting consortium, several *laboratories* or *data sources* generated the data, adding a hierarchical level above the *plate* and *batch* levels.

Cells in each well are treated with a specific *perturbation* (e.g., compound or gene overexpression). This provides *perturbation labels* to assess reproducibility across repeated measurements or *replicates*. We used several subsets of the JUMP-CP data, which we refer to as *datasets*. Aggregating perturbation features across all replicates in a dataset produces a *consensus profile*. For a small subset of compounds, we have drug *target labels*, i.e., proteins targeted by these drugs. Gene overexpression perturbations correspond to individual genes which can be annotated and grouped in *gene families*. To evaluate biological relevance, we used *target labels* for compounds and *gene family labels* for genetic perturbations.

### JUMP-CP training and validation sets

We used subsets of the JUMP Cell Painting dataset^27^ (cpg0016-jump) for training and evaluation of self-supervised learning (SSL) models. The complete JUMP-CP dataset (115 TB) includes 116,750 chemical perturbations, 12,602 gene overexpression and 7,975 CRISPR perturbations probed in a human cancer cell line (U2OS) in 5 replicates. Each chemical perturbation was screened by 5 out of the 10 consortium laboratories (“sources”) that used a standardized protocol but possibly different instrumentation. Genetic perturbations were screened solely by sources 4 and 7.

For model training, we used only images of cells treated with chemical perturbations. We used two training sets: a single-source and a multisource training set. The single-source training set consists of 391,815 images from JUMP-CP source 3, corresponding to 35,892 compounds with 1 replicate and 9 fields of view per well. The multisource training set contains 564,272 images from 4 JUMP-CP sources: source 2, source 3, source 6, and source 8. The multisource training set includes 5 replicates of 10,057 compounds from Selleckchem and MedChemExpress bioactive libraries, with two replicates originating from source 3. An overview of the JUMP-CP batches and plates used for model training is provided in **Supplementary Data 1**.

To assess our models, we used JUMP Target2 plates^27^ that contain 306 compounds with drug target labels. These plates were imaged in every experimental batch, enabling us to not only assess model performance using biological labels, but also evaluate batch and laboratory effects. Single-source (6 batches, 33,962 images) and multisource (16 batches, 75,545 images) validation sets were constructed using JUMP-Target2 plates from the respective sources of the single-source and multisource training sets. JUMP-CP batches and plates used for evaluation of models are listed in **Supplementary Data 2**.

### JUMP-CP gene overexpression test set

The JUMP-CP^27^ gene overexpression data (source 4) with Open Reading Frames (ORFs) was used for the final assessment of SSL models. A subset of the gene overexpression data was constructed by selecting ORF perturbations with high replicate correlations (𝑟 > 0.4) in the CellProfiler feature space, resulting in 5,198 ORFs. Gene group memberships were assigned to each ORF perturbation using the HUGO Gene Nomenclature Committee (HGNC) gene annotation (hgnc_complete_set_2022-10-01.txt). To ensure robust evaluation based on gene annotations, we selected only those gene groups with at least 4 unique ORFs. 1,970 ORF perturbations satisfied this criterion. The complete list of batches and plates is provided in **Supplementary Data 2**.

### Image preprocessing

The JUMP-CP consortium^27^ generated Cell Painting images with 5 color channels (Mito, AGP, RNA, ER, DNA) that were stored as individual TIFF files. To optimize data loading, we combined the single-channel images into 5-channel TIFF files, resulting in a 6-fold acceleration in training time. Prior to storage, we preprocessed the images: for each channel, intensities were clipped at 0.01^st^ and 99.9^th^ percentiles and scaled to the range [0,1]. Additionally, we calculated the Otsu threshold in the DNA staining channel and saved it as image metadata. During training, this threshold value enabled us to sample non-empty crops based on the minimum percentage of foreground area in the DNA channel.

### Augmentations for multichannel images

In the augmentation study, we trained 15 SimCLR models with the ResNet-50 backbone to test different image augmentation strategies. All models were trained and evaluated on the single-source data. We probed pairwise combinations of 5 augmentations: ‘Resize’, ‘Color’, ‘Drop channel’, ‘Gaussian noise’ and ‘Gaussian blur’. The ‘Flip’ augmentation that rotates an image by 180 degrees along the horizontal or vertical axes was used by default. ‘Resize’ generates crops with dimensions varying between 12% and 47% of the whole microscopy image and rescales the output to 224×224 pixels. The ‘Color’ augmentation consists of a random intensity shift: 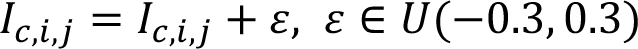 and a random brightness change: 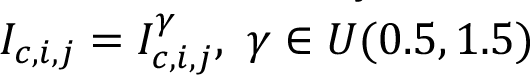 with intensity values restricted to [0, 1]. ‘Drop channel’ omits one channel from the image at random with probability 𝑝 = 0.5. The dropped channel is padded with zeros. ‘Gaussian noise’ adds random noise to the image: 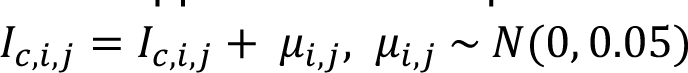. ‘Gaussian blur’ applies a Gaussian filter with a kernel size of 23 pixels and a standard deviation uniformly sampled from [0.1, 2]. Based on the results of the augmentation study, we applied only ‘Flip’ and ‘Color’ augmentations for training SimCLR and DINO. For training MAE, only the ‘Flip’ augmentation was used.

### Model training details

During training, we sampled random crops (224×224 pixels) from the images and provided these as inputs to the models. We only used image crops with cells (“cell-centered crops”), which was ensured by imposing a lower bound of 1% on the Otsu-thresholded area in the DNA channel. For SimCLR and DINO, we additionally applied the augmentation pipeline, described in “Augmentations for multichannel images”, to generate multiple views from the sampled crops. All input crops were centered and scaled using channel intensity means and standard deviations estimated over the entire training set. We used the small (ViT-S/16) and base (ViT-B/16) variants of the vision transformer, with a patch size of 16 pixels. The models with ViT-S/16 were trained for 200 epochs, while those with ViT-B/16 were trained for 400 epochs. We used the AdamW optimizer and saved checkpoints every 20 epochs. In addition to tracking the SSL training loss, which can be an unreliable indicator of downstream performance, we monitored training progress using the mean replicate correlation on the single-source evaluation set and selected the best-performing checkpoint for each model. A brief exposition of model-specific hyperparameters is provided below. For a comprehensive overview of training hyperparameters refer to **Supplementary Table 1**.

*DINO* was trained with a minibatch size of 128 (192 for ViT-B), a learning rate of 2 · 10^−3^ (1.5 · 10^&3^ for ViT-B), and a weight decay linearly increasing from 0.04 to 0.4. The learning rate followed a 20-epoch linear warmup followed by a cosine decay. For each image, 8 local crops (96×96) and 2 global crops (224×224) were sampled. DINO is a joint-embedding model with a student-teacher architecture^16^. DINO projects representations into a high-dimensional (here 20,000-dimensional) space where the temperature-scaled cross-entropy loss is optimized using a temperature of 0.1 for the student and 0.04 for the teacher network. The teacher temperature followed a linear warmup starting from 0.01 for 30 epochs.

*Masked autoencoder (MAE)* was trained with a minibatch size of 1024 (1536 for ViT-B), a learning rate of 6 · 10^−^( (9 · 10^−^ for ViT-B), and a weight decay of 0.05. The learning rate followed a 30-epoch linear warmup followed by a cosine decay. Given a partially masked input image, MAE reconstructs the missing regions using an asymmetric encoder-decoder architecture^17^, with a significantly smaller decoder. To accelerate data loading, 4 random crops were sampled from each image during training. The masking ratio was set to 50%, and image augmentation was performed using only horizontal and vertical flips.

*SimCLR* was trained with a minibatch size of 256, a learning rate of 1 · 10^−3^, and a weight decay of 0.1. The learning rate followed a 30-epoch linear warmup followed by a cosine decay.

SimCLR is a contrastive approach^15^ that aims to match augmented views from the same image in the representation space (“positives”), while pushing away representations from different images (“negatives”). The temperature-scaled cross-entropy loss was used as the objective function with a constant temperature value of 0.2.

### SSL inference and postprocessing

A feature extraction model maps an input image 𝐼 ∈ 𝑅^𝐶×𝐻×𝖶^ to a *d*-dimensional feature space through a mapping function 𝑓: 𝑅^𝐶×𝐻×𝖶^ → 𝑅^𝑑^. In DINO, MAE, and SimCLR, the mapping 𝑓(·) is performed by a vision transformer (ViT) backbone. The dimensionality 𝑑 of learned features depends on the network architecture, with 𝑑 = 384 for ViT-S and 𝑑 = 768 for ViT-B.

At inference, each microscopy image 𝐼, corresponding to a single field of view (FOV), was split into 224×224 image crops {𝑥_i_, 𝑖 = 1, … , 𝑁_𝑐𝑟𝑜𝑝𝑠_}. All image crops were passed through a pretrained ViT backbone to generate crop features 𝑓(𝑥_i_). Crops with no cells were excluded following the same criteria used during training. The resulting image crop features were aggregated using the arithmetic mean: 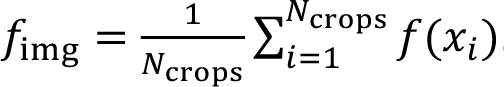. Well features were obtained by taking the mean across all FOV images: 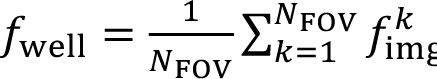.

We tested several feature postprocessing methods (**Extended Data Fig. 1-2**) and chose “sphering + MAD robustize” for SSL features. First, we removed well features with variance less than 1 · 10^−5^. We then applied a sphering transformation^39^ Φ to the remaining well features, followed by normalization using whole-plate median (𝑚𝑒𝑑) and median absolute deviation (𝑀𝐴𝐷):

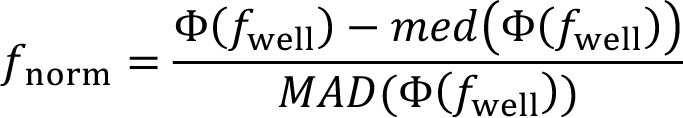

To generate perturbation profiles for downstream analyses, we averaged normalized well features 𝑓_*norm*_ across multiple replicates. Aggregation was performed at several levels: *batch-aggregated profiles* average all replicates within an experimental batch, *source-aggregated profiles* average all replicates within a JUMP-CP data source, and *consensus profiles* average all replicates within an entire dataset (single-source/multisource/gene overexpression).

### CellProfiler features

We used CellProfiler features provided by the JUMP-CP consortium (https://registry.opendata.aws/cellpainting-gallery/, for details see ^27^). CellProfiler features were normalized using whole-plate median and MAD (“MAD robustize” normalization), which was the best postprocessing method for CellProfiler features (**Extended Data Fig. 1e**). We tested several feature selection approaches (**Extended Data Fig. 2**) and selected the set of 560 features from the CPJUMP1 study^66^, in which low-variance and redundant features were removed based on a dataset with chemical and genetic (ORF and CRISPR) perturbations.

### Transfer learning

A vision transformer^35^ (ViT-S/16 or ViT-B/16) pretrained on the image classification task on ImageNet-1K was used to extract ‘transfer learning’ features. Each of the 5 channels was duplicated 3 times to generate pseudo-RGB images, which were individually passed through a pretrained ViT. The transfer learning features were obtained by concatenating individual channel features, resulting in 5 · 384 = 1920-dimensional feature vectors. As for SSL methods, low variance features (< 1 · 10^&<^) were removed before normalization. The transfer learning features were normalized using whole-plate median and MAD (“MAD robustize” normalization), which was the best postprocessing method for transfer learning (**Extended Data Fig. 1d**).

### Evaluation of reproducibility and biological relevance

We evaluated all features based on two key criteria: *reproducibility* using perturbation labels and *biological relevance* using drug target or gene family labels (see “Technical terminology”). To assess the sensitivity and precision of inferring ground-truth labels based on pairwise feature distances *D*(*f_i_, f_j_*), we followed an approach similar to ^66^. For each perturbation 𝑖, we define a neighborhood *N_i,d_* = {|*D*(*f_i_,f_i_*) ≤ *d*} consisting of all other perturbations 𝑗 within a cosine distance threshold 𝑑 of perturbation 𝑖. We then compared the label 𝑦_i_ of perturbation 𝑖 with the labels 𝑦_*j*_ of its nearest neighbors { 𝑗 ∈ 𝑁_i,𝑑_}. The precision 𝑃_i,𝑑_ and recall 𝑅_i,𝑑_ of matching labels for perturbation 𝑖 at distance threshold 𝑑 were calculated as:

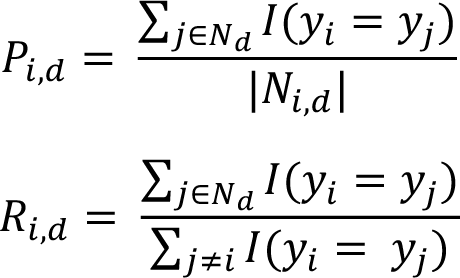

where 𝐼 is an indicator function, and |𝑁_i,𝑑_| is the size of the neighborhood of perturbation 𝑖.

Average precision (𝐴𝑃_i_) was computed for each perturbation by varying the distance threshold 𝑑 of the neighborhood 𝑁_i,𝑑_:

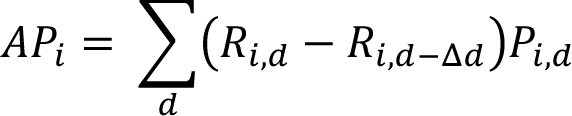

*Mean average precision (mAP)* was then calculated by averaging the AP values across all perturbations:

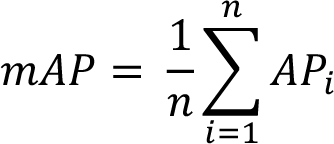

*Perturbation mAP*, measuring reproducibility, was estimated on batch-aggregated profiles (see “SSL inference and postprocessing”), thus quantifying the ability to match perturbations across batches. *Target mAP* was estimated on consensus profiles (see “SSL inference and postprocessing”) using drug target labels, focusing on biological content after technical variations were averaged out. For genetic perturbations, biological relevance was estimated using *gene family mAP*, calculated on consensus profiles with gene family labels. To evaluate the cross-source matching ability of features (**Extended Data Fig. 4e**), *perturbation mAP* was calculated on source-aggregated profiles. Along with AP values, F1-scores at *k* nearest neighbors were computed and visualized (**Extended Data Fig. 3**) to investigate whether some features worked better in specific *k* ranges.

The second class of metrics, widely used in morphological profiling^39,53,67^, reports the nearest neighbor (NN) accuracy estimated on well profiles with restrictions on the possible match. To evaluate reproducibility, the *not-same-batch (NSB) accuracy* restricts true positive matches to well profiles from different experimental batches. The *not-same-batch-or-perturbation (NSBP) accuracy* restricts true positive matches to profiles from both different batches and distinct perturbations. We used *perturbation NSB accuracy* to evaluate feature reproducibility across batches and *target NSBP accuracy* to evaluate biological relevance.

### UMAP embeddings

To visualize features in 2 dimensions, we generated UMAP (Uniform Manifold Approximation and Projection)^54^ embeddings using the first 200 principal components as input, with the correlation distance as the metric. For optimal visualization, we set the number of nearest neighbors to 50 and the minimum distance between points to 0.7 in the UMAP algorithm.

We selected the 20 drug targets with the highest mean F1-scores from the JUMP-Target2 plate^27^ annotation set. The F1-scores were determined by performing target classification using CellProfiler and DINO features. **Supplementary Table 2** provides the list of these 20 targets and their F1-scores stratified by feature type.

### Quantification of technical biases

The impact of technical variation was assessed by examining well profiles of the multisource validation dataset, which contained 24 replicates of each perturbation (see “Technical terminology”). To quantify batch and source effects for each feature type, we compared within- and between-cluster similarity and connectivity, using batch and source information as cluster labels. We used 3 metrics: *Silhouette scores*^68^, *Graph Connectivity (GC)*^69^ and *Local Inverse Simpson’s Index (LISI)*^69^.

The silhouette score measures the similarity of an observation 𝑖 to its own cluster (batch/source) relative to the nearest cluster^68^. It calculates the relative difference between the mean intra-cluster distance 𝑎(𝑖) and the mean nearest-cluster distance 𝑏(𝑖):

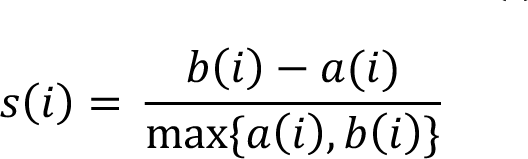

The silhouette score ranges from -1 to 1, with higher values indicating the observation is well matched to its own cluster and poorly matched to neighboring clusters. We compared distributions of silhouette scores for all well profiles clustered by batch or source (**Extended Data Fig. 6c,f**).

The *GC* and *LISI* metrics are based on a *k*-nearest neighbor (kNN) graph *G(V, E)*. This graph consists of vertices *V* corresponding to well profiles. Each vertex is connected to its *k* nearest neighbors based on pairwise cosine distances defining the edge set *E*. Let 𝐶 be a set of clusters, such as batches or sources. Taking only the vertices of a specific cluster 𝑐 ∈ 𝐶 induces a subgraph 𝐺_𝑐_(𝑉_𝑐_, 𝐸_𝑐_). GC measures the ratio between the number of vertices in the largest connected component *(LCC*) of 𝐺_𝑐_ and the total number of vertices in 𝐺_𝑐_, averaged across all clusters:

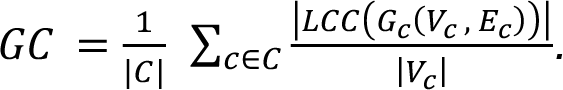

If the LCC of 𝐺_𝑐_is almost as large as 𝐺_𝑐_itself, this indicates that vertices from the same cluster are close together – a sign of batch/source effects. We reported GC for *k* = 1, 2, 3, 5, 10, 15 (**Extended Data Fig. 6a,d**).

LISI quantifies neighborhood diversity in the kNN-graph *G* using the inverse Simpson’s index:

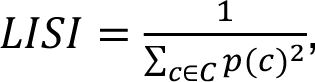

where 𝑝(𝑐) is the relative abundance of cluster 𝑐. LISI can be interpreted as the expected number of profiles to be sampled before two are drawn from the same cluster^61^. Higher *LISI* implies more diverse neighborhoods and lower batch/source effects. LISI was calculated for *k* = 15, 30, 60, 90 (**Extended Data Fig. 6b,e**). For both *GC* and *LISI*, the values for *k* are based on^69^.

For the sake of interpretability, we incorporated two baselines: 1) a Gaussian baseline, which simulates non-overlapping Gaussian (𝜎 = 1) clusters with the number of clusters equal to the number of batches/sources and with feature dimensionality identical to that of CellProfiler features; 2) a random baseline, which corresponds to the Gaussian baseline but with randomized cluster assignments.

### Hierarchical clustering of genetic perturbations

For the clustering analysis, we only considered HGNC gene families (see “Gene overexpression data”) that contained between 4 and 10 unique ORF perturbations. Since many of the gene families were heterogeneous and uncorrelated, we only selected the top 20 gene families with the highest within-family correlations in the respective feature space, resulting in 2 gene sets for DINO and CellProfiler (**Supplementary Data 3**).

To cluster these gene sets, we calculated the gene-gene correlation matrix within the respective representation space, which was then provided as input for hierarchical clustering using the complete linkage method and the Euclidean distance metric. To highlight biologically meaningful clusters in box frames, adjacent gene groups with at least 3 perturbations were identified visually and labeled with the majority gene family label.

To evaluate the quality of hierarchical clustering of genetic perturbations, we used *adjusted mutual information (AMI)*. For a given number of clusters, mutual information (MI) quantifies the dependence between cluster assignment labels 𝑋 and gene family labels 𝑌:

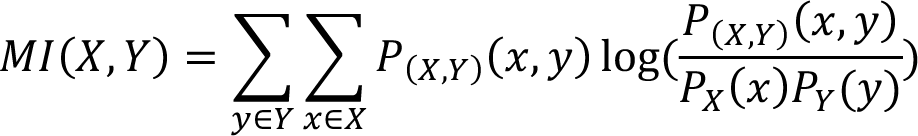

Adjusting mutual information (MI) for random chance results in AMI:

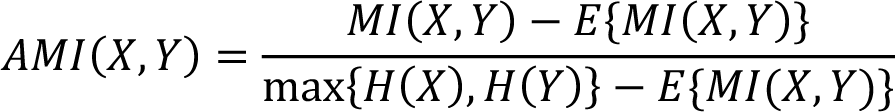

### Assessing the performance gap between supervised and self-supervised learning

We used DINO pretrained on the multisource JUMP-CP dataset to extract morphological features from Cell Painting images from a dataset of 30,000 small-molecule perturbations^70^. For bioactivity prediction, we only used 10,000 compounds with activity labels from a study^26^, in which convolutional neural networks (CNNs) were trained to predict compound activity across 209 ChEMBL assays. We trained a 3-layer fully connected neural network (FNN) on the extracted DINO features to predict compound activity. To ensure comparability with the CNNs trained on Cell Painting images directly, we used the same code base (https://github.com/ml-jku/hti-cnn), activity labels and train/validation/test splits as ^26^.

We included 6 CNNs from ^26^ as fully supervised baselines for bioactivity prediction: GapNet, ResNet, DenseNet, MIL-Net, M-CNN and SC-CNN. All CNNs, except Single-Cell CNN (SC-CNN), were trained end-to-end on Cell Painting images without segmentation. A 3-layer FNN trained on CellProfiler features was incorporated as an additional baseline from ^26^. The results for the CNNs and CellProfiler FNN were taken from the original publication. For details on the CNN and FNN architectures and training methodology, refer to ^26^.

Following ^26^, we evaluated the performance of the FNN trained on DINO features using area under the receiver operating characteristic curve (AUCROC) as the primary metric. The AUCROC values for each assay were obtained by averaging across 3 test splits. The mean AUCROC across 209 assays and the standard deviation are reported in **Extended Data Table 1**. Similarly, F1-scores were computed and the mean and standard deviation across all assays are also provided. To illustrate the performance gap across different data availability regimes, we grouped the assays into 5 bins based on the number of activity labels: [< 50, 50-100, 100-500, 500-1000, 1000-3000]. AUCROC values were visualized for these 5 assay groups in **Extended Data Figure 9**.

### Implementation details

All self-supervised learning (SSL) models were implemented in Python 3.9.7 using PyTorch^71^ v1.10.2 and PyTorch Lightning v1.6.3. SSL model training and inference were conducted on NVIDIA Tesla V100 GPUs with a VRAM of 32GB. Feature postprocessing (sphering, MAD robustize, standardize) was carried out using pycytominer v0.2.0. Mean average precision (mAP), NSB and NSBP accuracies were computed using custom Python functions, with average precision (AP) and accuracy scores computed using the scikit-learn^72^ v1.0.2 implementation. PCA and UMAP were performed using scikit-learn v1.0.2 and umap-learn v0.5.3. The silhouette scores, kNN graphs, and the simulated Gaussian baseline for batch and source effect quantification were computed using scikit-learn v1.0.2. The LISI scores were calculated using HarmonyPy^61^ v0.0.9. Adjusted mutual information (AMI) of gene family clustering was calculated using scikit-learn v1.0.2. The visualization of UMAP was performed using matplotlib v3.5.0 and seaborn v0.11.2. The visualization of evaluation metrics and hierarchical clustering of ORF perturbations was performed in R 4.1.2.

### Software availability

The code for training, inference, and evaluation of the self-supervised learning (SSL) models used in this study is provided in **Supplementary Code** and will be made publicly available on GitHub upon publication. The code is distributed under the BSD 3-Clause License. The model weights are provided and intended for non-commercial use only.

## Acknowledgements

We would like to express our gratitude to the following individuals and organizations for their contributions to this work. We thank Bayer AG, which enabled the successful completion of this work through the Life Science Collaboration (LSC) project ’Picasso’. We thank all members of the JUMP-CP consortium for producing the Cell Painting dataset that was used for training SSL models in this study. We would like to thank Djork-Arne Clevert for his support and guidance throughout this project. Finally, we thank all members of the Machine Learning Research group at Bayer AG for their kind advice and support.

## Author contributions

VK and PAMZ designed the study. PAMZ supervised the study. VK prepared training and evaluation data. VK and NA implemented the self-supervised learning (SSL) framework including data loading, augmentation, and training pipelines. VK and NA trained SSL models. MO and VK designed and implemented a cloud-based inference pipeline. VK and MO produced representations for the gene overexpression data using the inference pipeline. VK conducted an augmentation study. VK and NA performed evaluations of SSL representations. FM conducted batch and laboratory effect analysis on the SSL representations. MH trained a bioactivity prediction model on DINO features and compared it with supervised CNNs. VK designed the figures, with contributions from PAMZ, MO, FM and NA. VK, NA and PAMZ wrote the manuscript with inputs from MO, FM, TK and DG. All authors reviewed and approved the final version.

## Competing interests

VK, MO, FM, MH, TK, DG, and PAMZ are employees of Bayer AG.

## Extended Data Figures

**Extended Data Figure 1:**
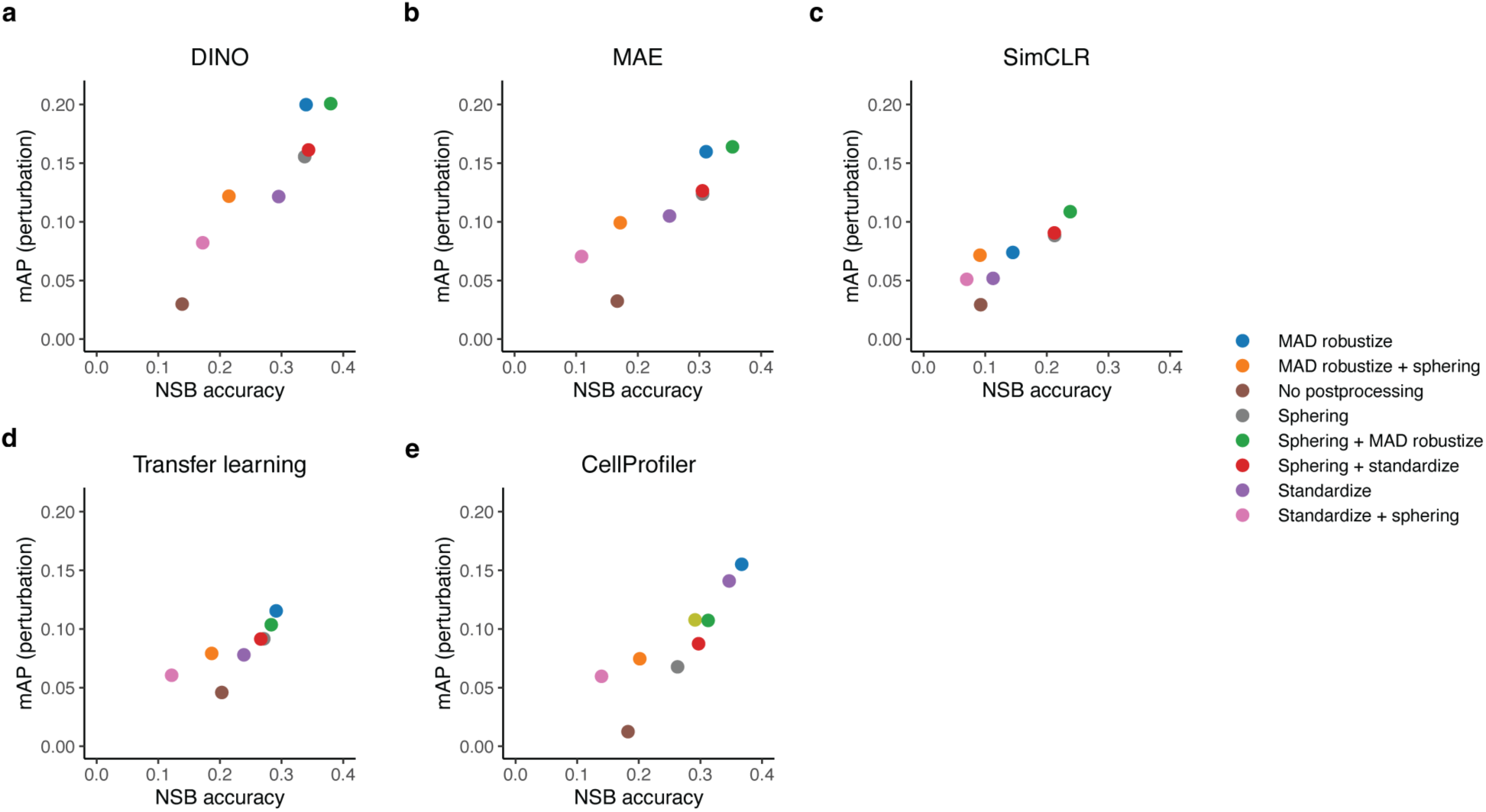
Comparison of postprocessing methods for various representations. Impact of normalization methods on reproducibility metrics *not-same-batch (NSB) accuracy* and *perturbation mean average precision (mAP)* computed on the multisource evaluation set. SSL models were trained on the multisource training set with the ViT-S architecture. Pairwise combinations of 3 normalization methods were tested. MAD robustize: center and scale each feature using plate median and median absolute deviation (MAD). Standardize: center and scale each feature using DMSO negative control mean and standard deviation. Sphering: center and scale using DMSO negative control mean and standard deviation and transform the data using the eigenvector matrix of the DMSO covariance matrix.

**Extended Data Figure 2:**
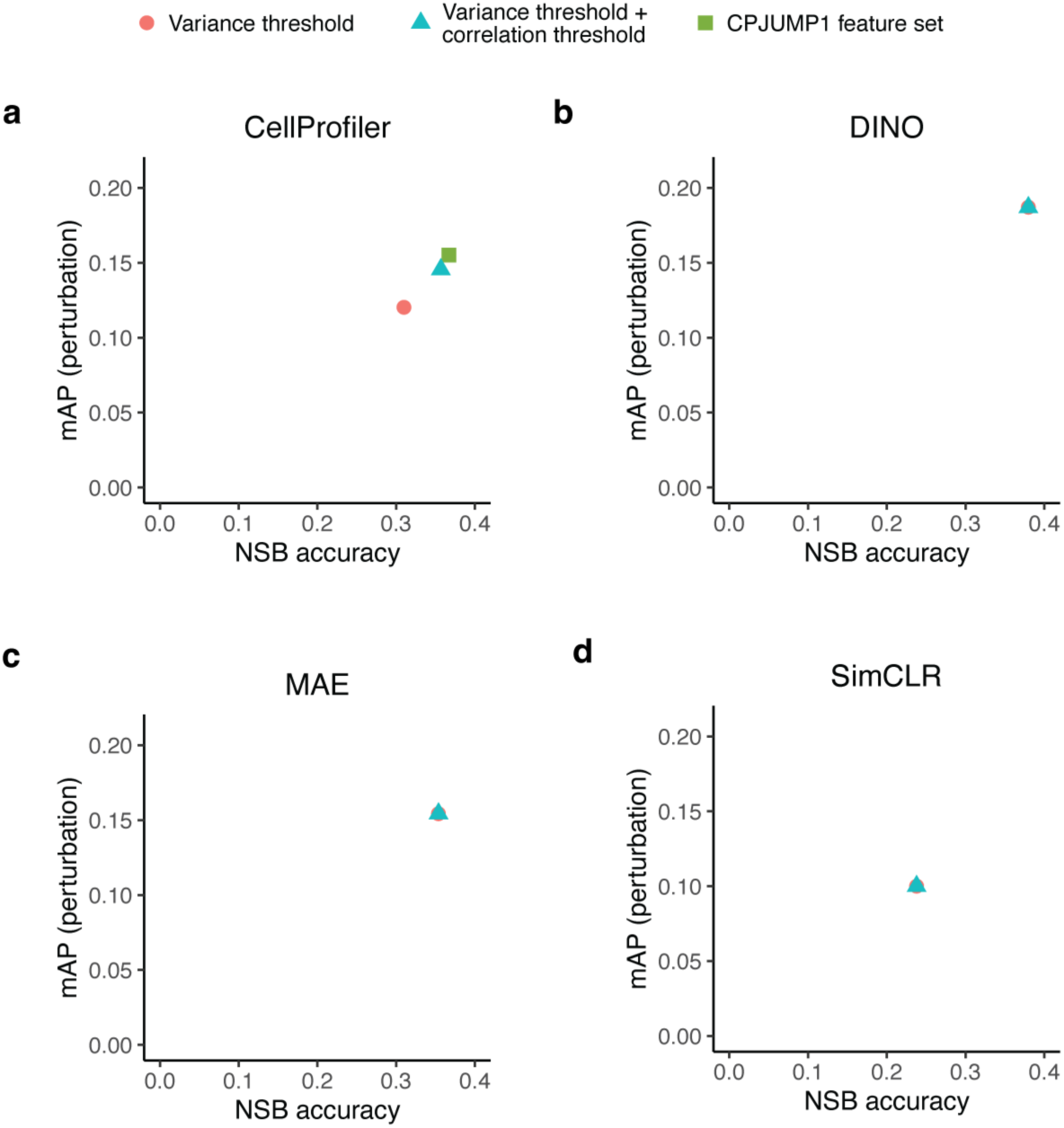
Comparison of feature selection methods for various representations. Impact of feature selection methods on reproducibility metrics *not-same-batch (NSB) accuracy* and *perturbation mean average precision (mAP)* computed on the multisource evaluation set. SSL models were trained on the multisource training set with the ViT-S architecture. Two normalization methods were tested: Variance threshold was used to remove low-variance features, and variance threshold + correlation threshold was used to further eliminate redundant features. For CellProfiler, the CPJUMP1 feature set was also included, selected based on the CPJUMP1 dataset^1^, which comprises chemical, ORF, and CRISPR perturbations.

**Extended Data Figure 3:**
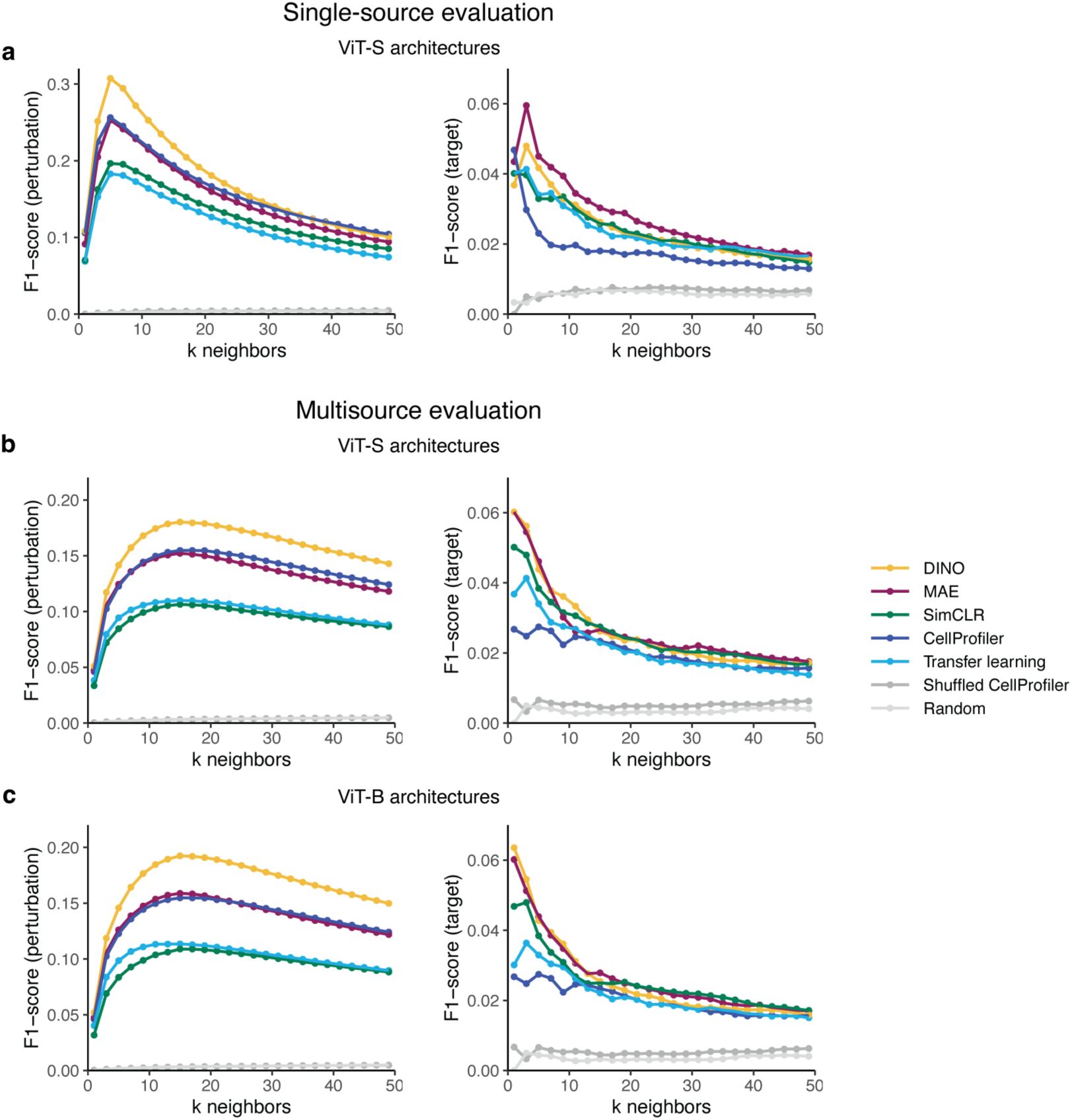
F1-scores of SSL models on single-source and multisource evaluation sets. F1-score curves for matching perturbation (left) and target (right) labels for a range of nearest neighbors *k.* SSL models were trained on the multisource training set. Colors indicate different models and two randomized baselines: Shuffled CellProfiler (CellProfiler features with shuffled labels) and Random (random normally distributed features). a) Performance of ViT-S architectures on the single-source evaluation set. b) Performance of ViT-S architectures on the multisource evaluation set. c) Performance of ViT-B architectures on the multisource evaluation set.

**Extended Data Figure 4:**
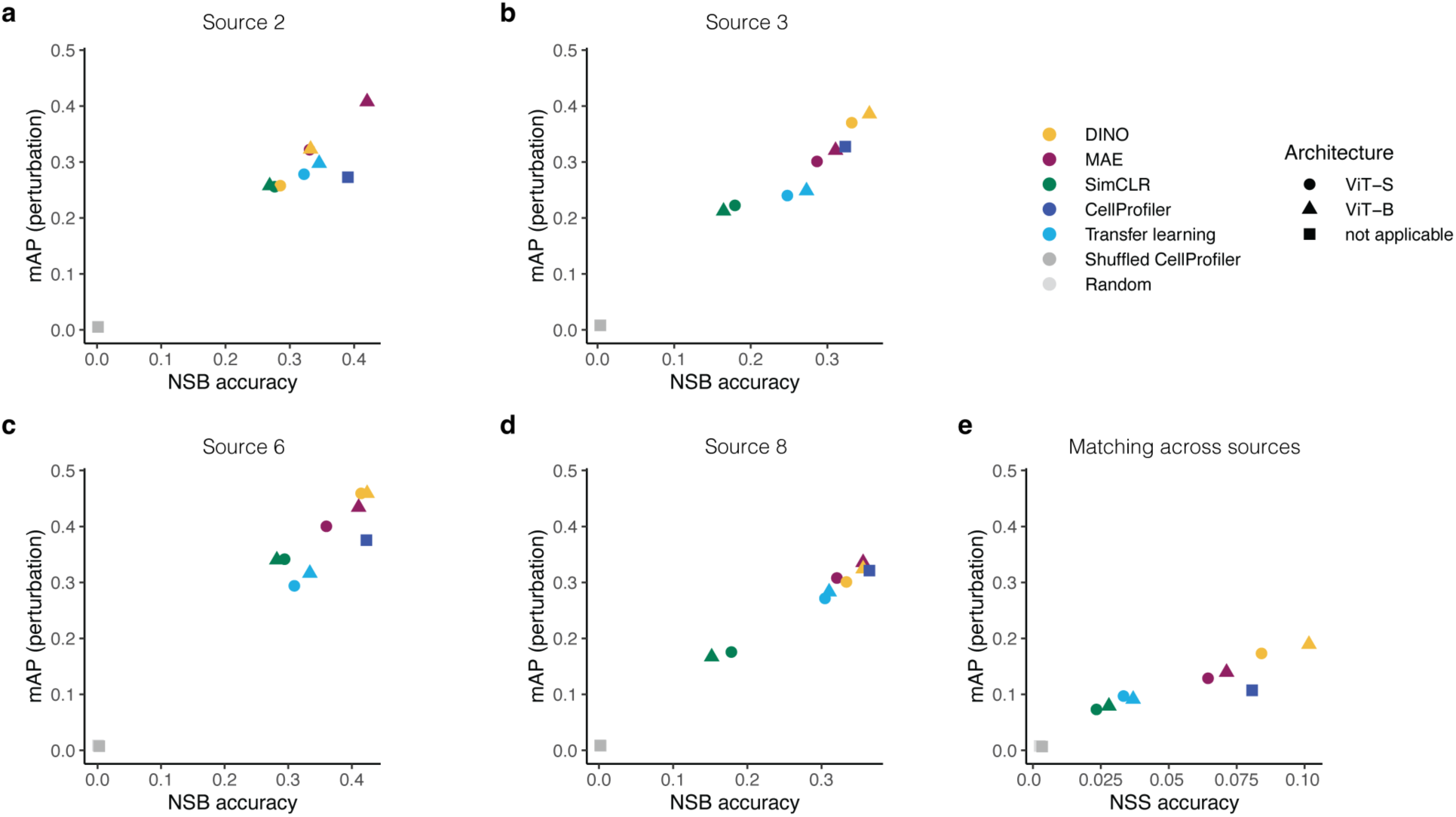
Evaluation of SSL methods for individual JUMP-CP data sources. Perturbation reproducibility metrics computed on the multisource evaluation set for 4 JUMP-CP consortium data sources. SSL models were trained on the multisource training set. Shapes indicate the ViT architecture and colors indicate the model. Two randomized baselines were included: Shuffled CellProfiler (CellProfiler features with shuffled labels) and Random (random normally distributed features). a)-d) Matching of perturbation labels for each individual JUMP-CP source. e) Matching of perturbation labels across all 4 sources. *Mean average precision (mAP)* was computed on source-aggregated profiles. *Not-same-source (NSS) accuracy* quantifies the accuracy of matching well profiles across different data sources using a nearest neighbor classifier.

**Extended Data Figure 5:**
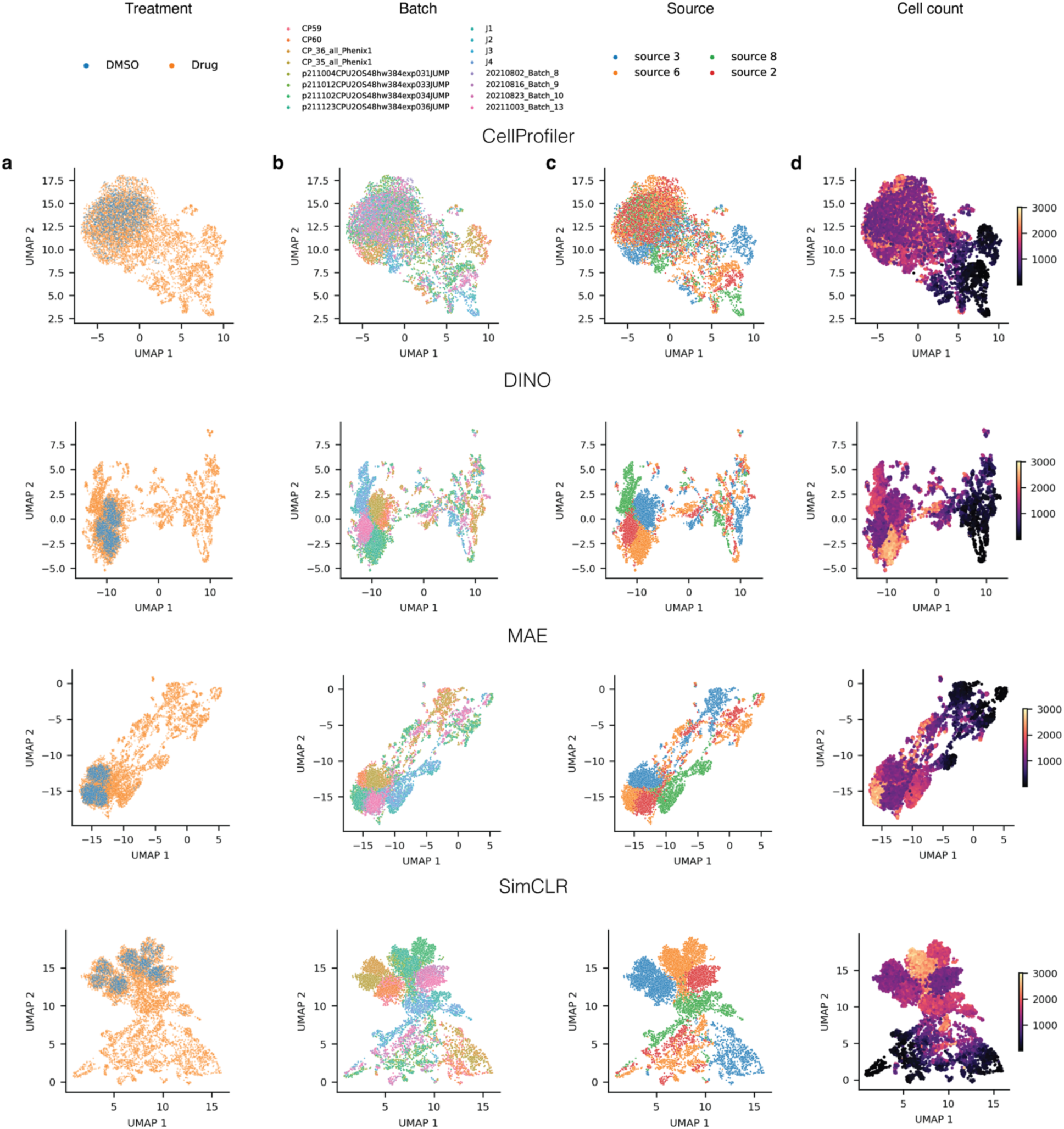
UMAP projections of the multisource evaluation set reveal batch and source effects. UMAP projections of well representations for SSL models and the CellProfiler baseline. All SSL models were trained on the multisource training set with the ViT-S architecture. The points, corresponding to wells, are colored by a) treatment (DMSO negative control vs drug), b) batch, c) source, and d) cell count.

**Extended Data Figure 6:**
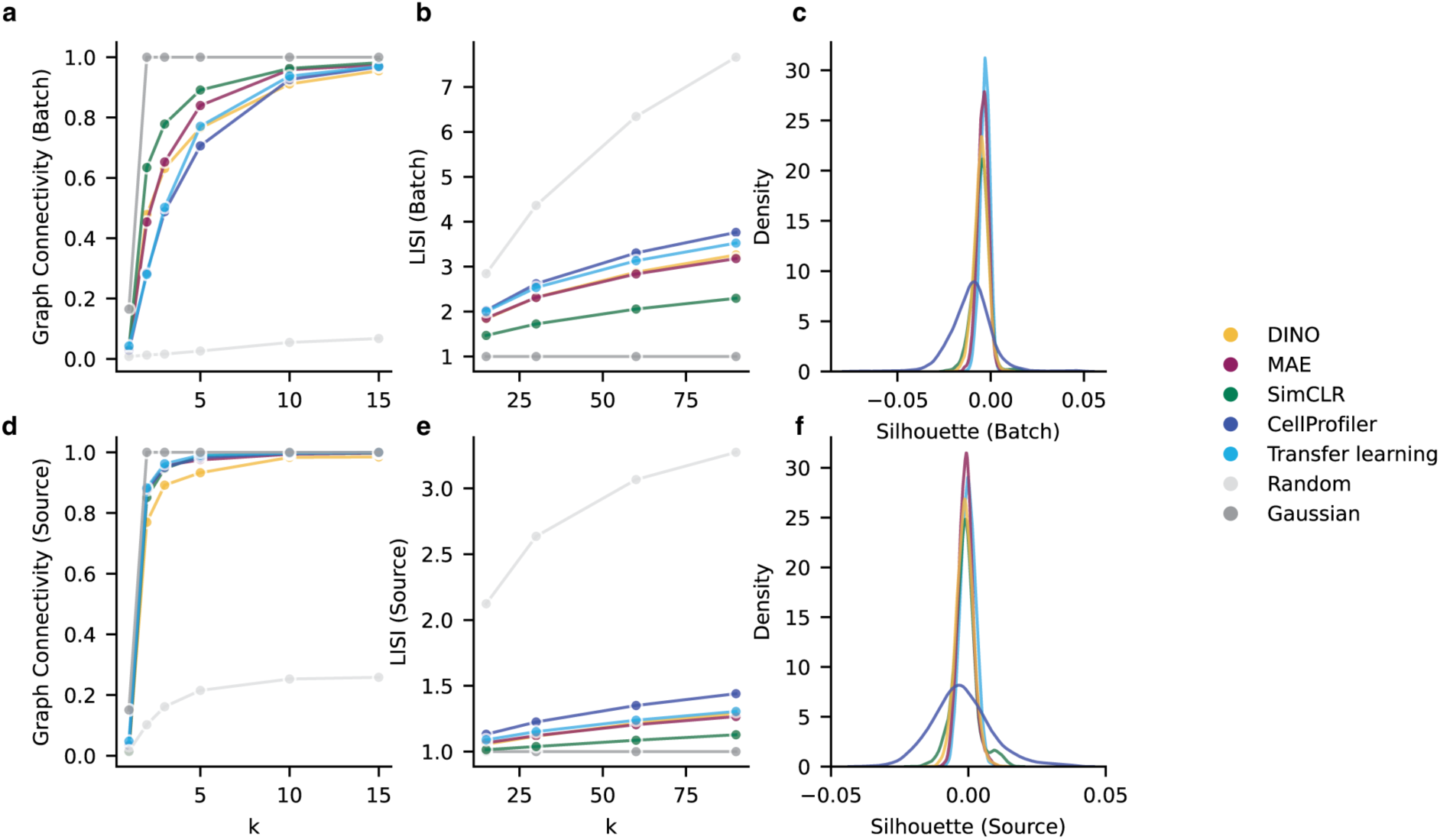
Quantitative assessment of batch and laboratory effects for SSL and baseline methods. Impact of technical variation on SSL and baseline representations assessed on the multisource evaluation set. SSL models were trained on the multisource training set with the ViT-S architecture. Batch and laboratory (source) effects were assessed using graph connectivity, Local Inverse Simpson’s Index (LISI), and silhouette scores. For reference, two synthetic representations were included: ‘Gaussian’ with non-overlapping Gaussian clusters and ‘Random’ with shuffled Gaussian cluster labels. a)-c) Metrics for batch effects. d)-f) Metrics for source effects.

**Extended Data Figure 7:**
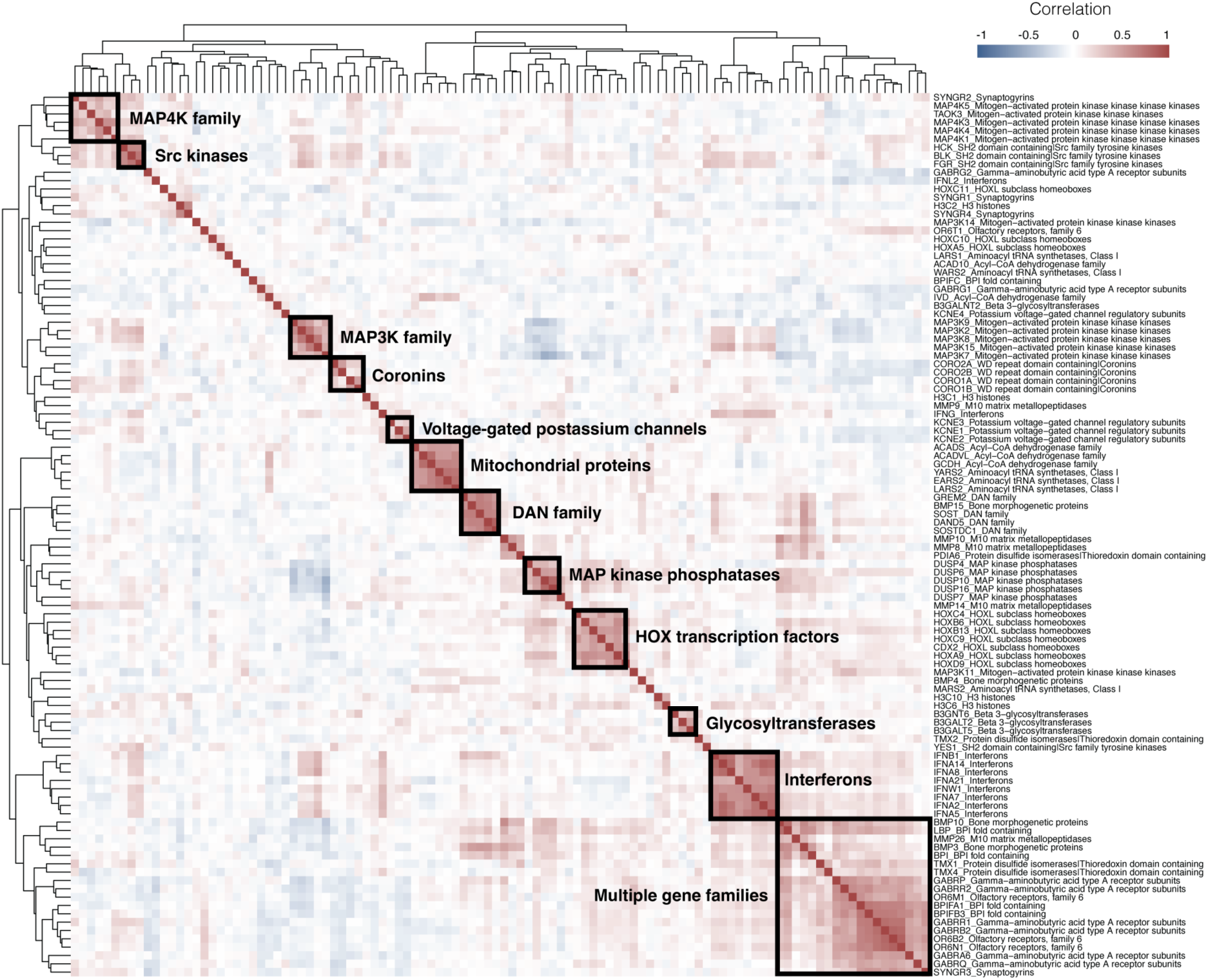
Hierarchical clustering of DINO representations for selected genetic perturbations. Hierarchical clustering of the pairwise correlation matrix of DINO representations for the subset of 20 gene families with the highest within-group correlations in the DINO feature space. Rows are labeled with perturbation and gene family names separated by an underscore (e.g. “HOXA9_HOXL subclass homeoboxes”). Clusters recapitulating gene groups, such as “HOX transcription factors” and “mitochondrial proteins”, are highlighted. DINO was trained on the multisource data with the ViT-S architecture.

**Extended Data Figure 8:**
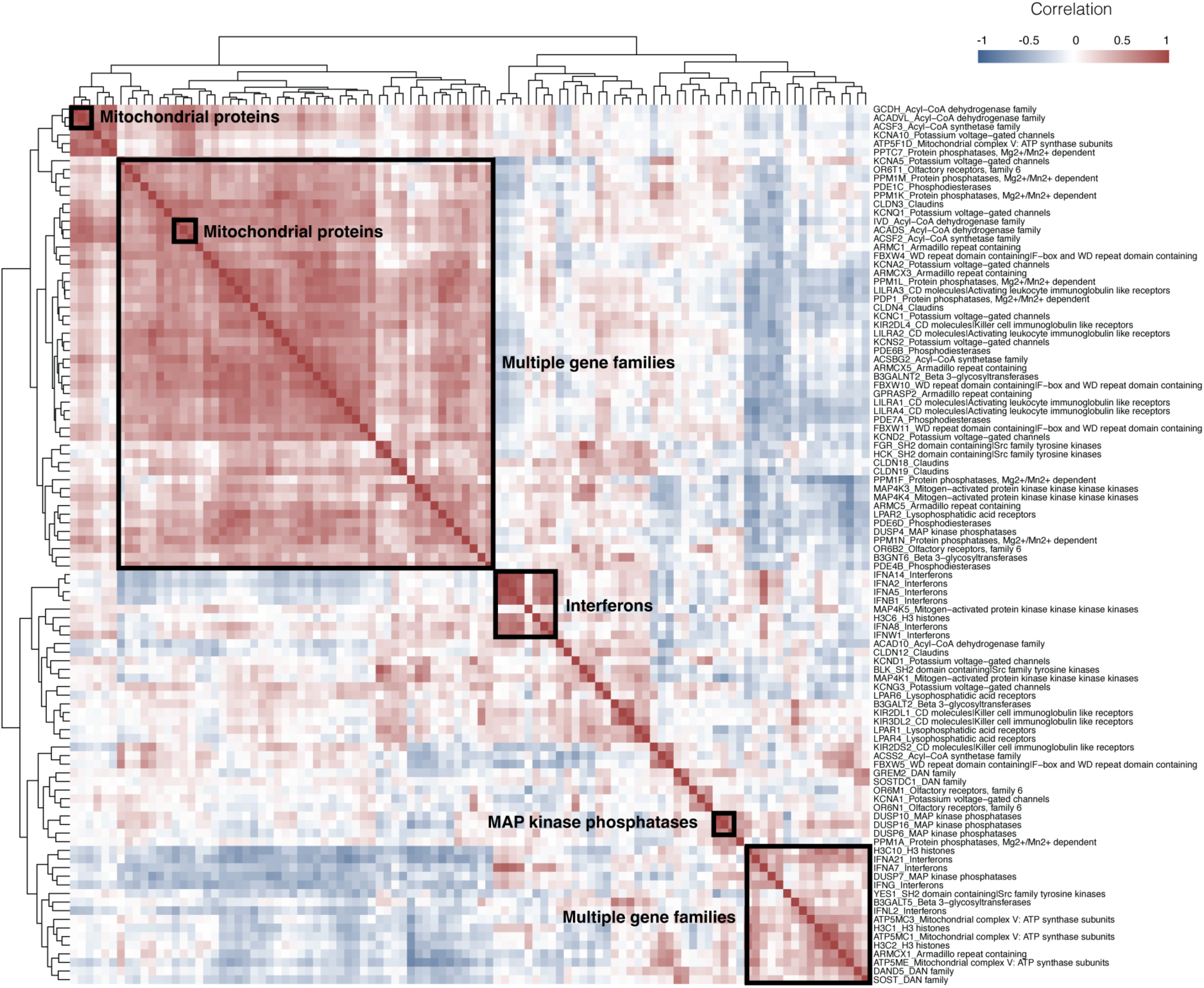
Hierarchical clustering of CellProfiler representations for selected genetic perturbations. Hierarchical clustering of the pairwise correlation matrix of CellProfiler representations for the subset of 20 gene families with the highest within-group correlations in the CellProfiler feature space. Rows are labeled with perturbation and gene family names separated by an underscore (e.g. “IFNA2_Interferons”). Clusters recapitulating gene groups, such as “mitochondrial proteins” and “MAP kinase phosphatases”, are highlighted.

**Extended Data Figure 9:**
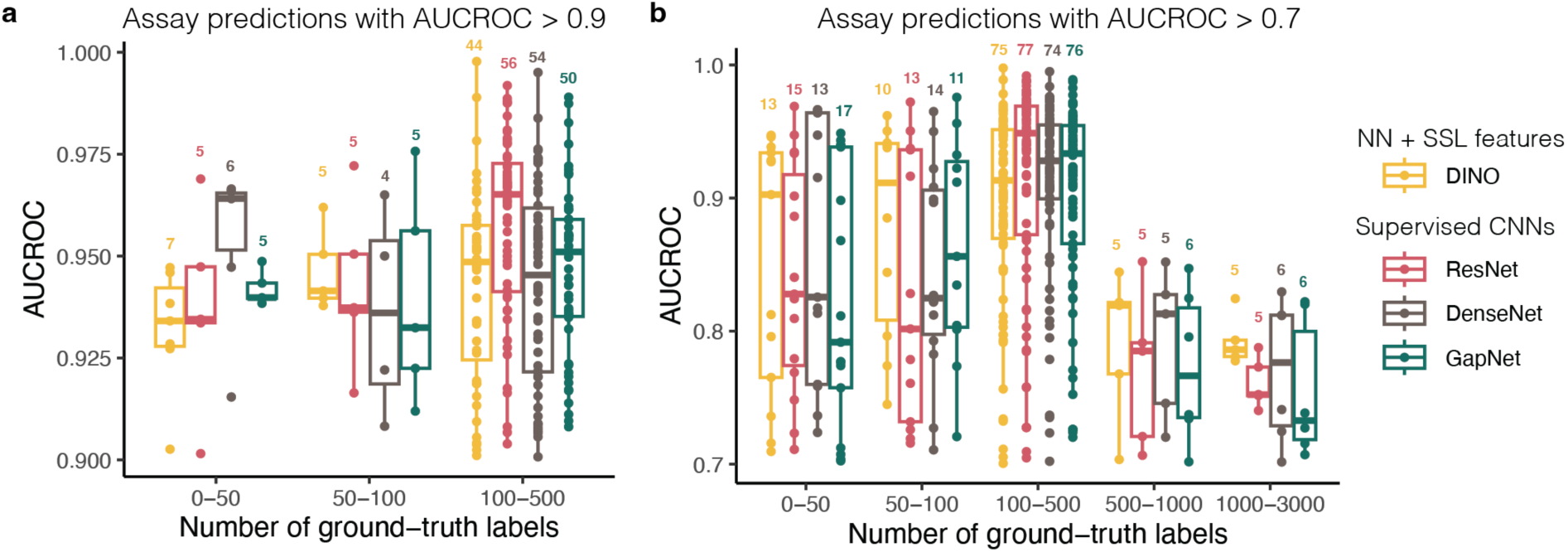
Performance of bioactivity prediction models stratified by assay data abundance. Compound activity prediction performance was compared between a neural network (NN) trained on DINO features and the top 3 convolutional neural networks (CNNs) from Hofmarcher et al.^2^ across 209 ChEMBL assays grouped by the number of available activity labels (x-axis). Models were evaluated by the area under the receiver operating characteristic curve (AUCROC, y-axis). Assay AUCROC values are plotted for each model, with median and upper/lower quartile values summarized in boxplots. Only assays with **a)** AUCROC > 0.9 and **b)** AUCROC > 0.7 were considered. The numbers above boxplots indicate the assay counts per group.

**Extended Data Figure 10:**
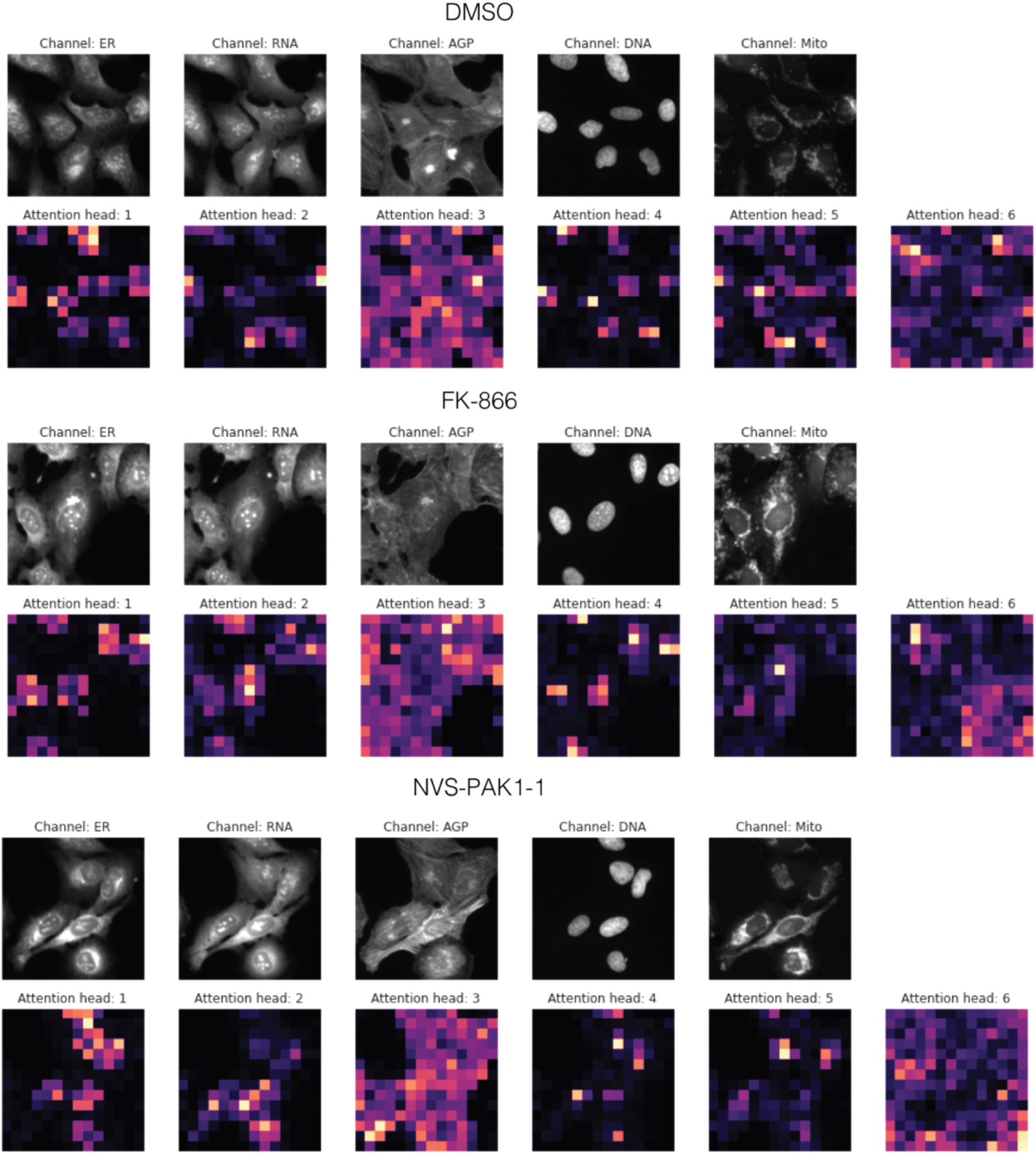
DINO self-attention maps. Cell Painting image crops and self-attention maps of the DINO attention heads in the last layer. Example images for DMSO, FK-866 and NVS-PAK1-1. The color scale in the self-attention maps represents the level of attention from the DINO [CLS] token, with lighter areas indicating higher attention. DINO was trained on the multisource data with the ViT-S architecture.

## Extended Data Tables

**Extended Data Table 1:** Performance comparison of bioactivity prediction models using Cell Painting. Eight deep learning methods were used to predict bioactivity labels for 209 ChEMBL assays using Cell Painting (see Methods). The methods included a neural network (NN) trained on DINO features, 6 convolutional neural networks (CNNs) trained directly on Cell Painting images, and an NN trained on CellProfiler features. Performance was evaluated on held-out test data, reporting means and standard deviations of AUCROC and F1-score values across all assays. Additionally, the number of assays with AUCROC above 0.9, 0.8, and 0.7 is reported for each method. The DINO results were obtained by applying a DINO model pretrained on the JUMP-CP data to the same dataset used in Hofmarcher et al.^3^ The remaining results are from Table 1 of Hofmarcher et al.^3^ Methods marked with ^†^ require cell-level segmentation, while the others use whole images as input. AUCROC: area under the receiver operating characteristic curve.

| Method | Input | Arch. | AUCROC | F1-score | AUC > 0.9 | AUC > 0.8 | AUC > 0.7 |
| --- | --- | --- | --- | --- | --- | --- | --- |
| ResNet | Images | CNN | 0.731 ± 0.19 | 0.508 ± 0.30 | 68 | 94 | 119 |
| DenseNet | Images | CNN | 0.730 ± 0.19 | 0.530 ± 0.30 | 61 | 98 | 121 |
| GapNet | Images | CNN | 0.725 ± 0.19 | 0.510 ± 0.29 | 63 | 94 | 117 |
| **DINO** | **SSL features** | NN | **0.723 ± 0.18** | **0.507 ± 0.31** | **56** | **84** | **108** |
| MIL-Net | Images | CNN | 0.711 ± 0.18 | 0.445 ± 0.32 | 61 | 81 | 105 |
| M-CNN | Images | CNN | 0.705 ± 0.19 | 0.482 ± 0.31 | 57 | 78 | 105 |
| SC-CNN<sup>†</sup> | Images | CNN | 0.705 ± 0.20 | 0.362 ± 0.29 | 61 | 83 | 109 |
| CellProfiler<sup>†</sup> | Handcrafted features | NN | 0.675 ± 0.20 | 0.361 ± 0.31 | 55 | 71 | 90 |

**Extended Data Table 2:** Inference speed and cloud cost comparison for DINO and CellProfiler. Comparison of processing speed and cloud costs between DINO and CellProfiler pipelines for the analysis of 28 plates in the AWS cloud. The cloud costs include only EC2 instance charges.

| Method | Average time per plate | Processing time for 28 plates | AWS EC2 instance type | Cloud cost per plate |
| --- | --- | --- | --- | --- |
| DINO | 1.3 min. | 36.6 min. | 12 x g4dn.xlarge | \$0.17 |
| CellProfiler | 66.7 min. | 1867.2 min. | 8 x r5.24xlarge + 4x r5.8xlarge | \$10 |

## Footnotes

1 Chandrasekaran, S. N. *et al.* Three million images and morphological profiles of cells treated with matched chemical and genetic perturbations. 2022.01.05.475090 Preprint at https://doi.org/10.1101/2022.01.05.475090 (2022).

2 Hofmarcher, M., Rumetshofer, E., Clevert, D.-A., Hochreiter, S. & Klambauer, G. Accurate Prediction of Biological Assays with High-Throughput Microscopy Images and Convolutional Networks. *J Chem Inf Model* 59, 1163–1171 (2019).

3 Hofmarcher, M., Rumetshofer, E., Clevert, D.-A., Hochreiter, S. & Klambauer, G. Accurate Prediction of Biological Assays with High-Throughput Microscopy Images and Convolutional Networks. *J Chem Inf Model* 59, 1163–1171 (2019).

## Notes

### Summary of Updates

Figure 5 has been updated. New results quantifying the performance gap between self-supervised and supervised learning for Cell Painting have been incorporated. The Results section has been streamlined, with technical details now moved to the Methods section. The Methods section has been expanded to enhance technical and mathematical clarity. The Introduction and Discussion have been revised to address potential misconceptions. The author list has been updated based on additional contributions.

